# Measuring rank robustness in scored protein interaction networks

**DOI:** 10.1101/502302

**Authors:** Lyuba V. Bozhilova, Alan V. Whitmore, Jonny Wray, Gesine Reinert, Charlotte M. Deane

## Abstract

**Background:** Protein interaction databases often provide confidence scores for each recorded interaction based on the available experimental evidence. Protein interaction networks (PINs) are then built by thresholding on these scores, so that only interactions of sufficiently high quality are included. These networks are used to identify biologically relevant motifs or nodes using metrics such as degree or betweenness centrality. This type of analysis can be sensitive to the choice of threshold. If a node metric is to be useful for extracting biological signal, it should induce similar node rankings across PINs obtained at different reasonable confidence score thresholds.

**Results:** We propose three measures—rank continuity, identifiability, and instability—to evaluate how robust a node metric is to changes in the score threshold. We apply our measures to twenty-five metrics and identify four as the most robust: the number of edges in the step-1 ego network, as well as the leave-one-out differences in average redundancy, average number of edges in the step-1 ego network, and natural connectivity. Our measures show good agreement across PINs from different species and data sources. Analysis of synthetically generated scored networks shows that robustness results are context-specific, and depend both on network topology and on how scores are placed across network edges.

**Conclusion:** Due to the uncertainty associated with protein interaction detection, and therefore network structure, for PIN analysis to be reproducible, it should yield similar results across different confidence score thresholds. We demonstrate that while certain node metrics are robust with respect to threshold choice, this is not always the case. Promisingly, our results suggest that there are some metrics that are robust across networks constructed from different databases, and different scoring procedures.

## Background

Protein interaction networks (PINs) are models of cellular architecture in which proteins are represented by nodes and the biologically meaningful interactions between them are represented by edges. PIN analysis has a wide range of applications in bioinformatics [1–3], and in particular in drug discovery [4–6]. For example, it can be used to predict protein function [7–10] and disease relevance [11–13], as well as to identify possible drug targets, especially in the case of multi-target drug discovery [14]. A common aim of PIN analysis is the identification of key actors in the network, e.g. for the purposes of drug target choice [15–17].

PINs can be built using both physical and/or functional interactions, which can in turn be obtained from a range of experimental and *in silico* techniques [18, 19]. There exist numerous databases of protein interactions which vary in data type, data collection methods, and content curation [20, 21]. High-throughput interaction data based on yeast two-hybrid assays [22], tandem affinity purification [23] or gene co-expression [24] are often subject to high false positive and false negative rates (e.g. [25–27]. Different approaches to data curation are used to correct for this. Some databases, such as STRING [28], HitPredict [29], IntAct [30], and HIPPIE [31], quantify the strength of the supporting evidence for each reported interaction by assigning a confidence score to it. These confidence scores are often a combination of different sub-scores (e.g. based on different evidence channels in STRING), each of which is calculated in a custom, source-specific way. The final combined scores are usually scaled between 0 and 1 but are not readily interpretable, and, as we illustrate, tend not to be comparable across databases. Such scores are designed to provide a comparison between different interactions (an interaction with confidence score of 0.40 is supported by weaker evidence than an interaction with score of 0.90), so researchers can control strength of evidence by imposing a score threshold. STRING, for example, suggests thresholds of 0.15 (low confidence), 0.40 (medium confidence), 0.70 (high confidence) or 0.90 (highest confidence), whereas HitPredict identifies all interactions scoring below 0.28 as medium-high confidence, and interactions scoring above as high confidence.

Due to the wide range of available resources and quality assessment tools for protein interaction data a number of different networks can be built to model the same part of the interactome. Even when the data type and source are fixed, a threshold on data quality is often chosen, either by database curators, or explicitly by researchers during network construction. Only interactions which meet that threshold contribute to the final network structure (see [28] for a discussion on scores and thresholding, and [32] for a particular example). Such data preprocessing choices play an important role in PIN structure and we show that they can have an effect on any further network analysis.

Given interaction detection error rates, and the incomplete coverage of interaction detection experiments [33, 34], it is extremely unlikely that any one PIN is a perfect representation of the underlying biological processes it aims to model, regardless of how it is constructed. The effect of error, or noise, on networks, such as missing or misplaced edges, is an established research topic in network science [35], and is often studied via network perturbation [36]. In the context of PINs, stochastic models of noise on network edges have been used to re-score interactions [9], determine optimal score thresholds [37], and have been incorporated into community detection [38, 39]. However, these models often rely on assumptions about the behaviour of the error: for example, they may assume that interactions are equally likely to be detected regardless of the properties of the proteins involved. Such assumptions do not necessarily hold in the context of protein interaction detection [40]. Moreover, any approach which incorporates interaction scores directly within PINs (e.g. by modelling them as weighted networks), implicitly relies both on the interpretability and the accuracy of the scores themselves, both of which can change over time and across data sources. The relative scarcity of high quality interaction data and lack of sufficiently good PIN-like random network models further complicate the validation of such approaches.

Rather than considering a single network, the observation of which is subject to a difficult to model noise process, assessing the robustness of a PIN analysis pipeline can be done by repeating the analysis across different networks representing the same part of the interactome. One way to do this would be to consider building different networks from the same scored interaction database by varying the confidence score threshold. We postulate that network features which are persistent across different thresholds are more likely to be informative of the biological state of interest than features which are only present at isolated thresholds. This hypothesis is in line with network research in neuroscience, where a similar heuristic is employed to identify which parts of a brain network are active across different observations [41].

In this paper we provide a framework for assessing the robustness of node metrics to threshold choice. Our framework is based on a measure of rank similarity described by [42]. We introduce three robustness measures—rank continuity, identifiability, and instability—which can be used to quantify how consistent a node metric is across different thresholds. Our methodology is particularly relevant to cases where a node metric is used to identify highly ranking nodes, which may correspond to key proteins in a particular process, for example for the purposes of drug target identification [16].

By analysing the effects of threshold change on a set of twenty-five node metrics across four scored PINs we show that some node metrics tend to be more robust—and are therefore possibly more relevant to biological research—than others. The node metrics studied include standard node centralities, such as degree and betweenness, as well as leave-one-out difference (LOUD) metrics, which measure the effect of isolating a node on global network summaries such as the global clustering coefficient. We identify the number of edges in the step-one ego network, and LOUD average redundancy [43], LOUD number of edges in the step-one ego network, and LOUD natural connectivity [44], as significantly more robust to threshold choice than more commonly used metrics, such as local clustering coefficient, betweenness centrality, and in some cases even degree.

Promisingly, our results show good agreement between networks from different organisms and databases. Complemented with analysis of synthetic data, we further show that robustness depends both on network topology and on score allocation across network edges.

## Results

### Thresholding effects

The reliability of a detected interaction between two proteins is often quantified by a confidence score, with lower scores corresponding to weaker interaction evidence. When PINs are constructed from such data, a threshold on the confidence scores is usually applied in an attempt to filter out spurious interactions. While it is possible to incorporate the confidence scores as edge weights in the network, these scores represent neither interaction strength, nor distance, making classical weighted network analysis techniques difficult to interpret. Moreover, confidence scores vary across databases, both in their derivation and interpretation, as well as in their values.

Confidence scores are designed specifically to allow researchers a degree of control over data quality, usually through thresholding. Threshold choice over the confidence scores introduces a trade-off between the numbers of false positive and false negative interactions. A low threshold may introduce many interactions which have been detected experimentally, but which have no biological relevance, while a high one will reduce the number of such false positives, but may also lead to more genuine interactions being excluded from the network.

Imposing different thresholds will affect PIN structure, and may affect PIN analysis in complex, and potentially difficult to predict, ways. Some metrics, such as edge density, node degree, and natural connectivity, will decrease monotonically with threshold increase. Other network metrics, such as clustering coefficients and betweenness centrality, do not necessarily behave monotonically and it is unclear how to predict their rate of change (or even its direction) between thresholds.

To examine the effect of threshold choice we considered three full organism networks obtained from the STRING database—*Plasmodium vivax* (PVX), *Escherichia coli* (ECOLI), and *Saccharomyces cerevisiae* (YEAST). STRING suggests using one of four thresholds as a default—low (0.15), medium (0.40), high (0.70), and highest (0.90). For each threshold, an unweighted network between the proteins is constructed which includes only those edges for which the score is at least as high as the threshold. The average degree in each of the three STRING networks analysed decreased with increasing thresholds, from over 200 at low confidence, to under 25 at highest confidence (Figure 1A). In the PVX network, the average local clustering decreased monotonically from 0.50 down to 0.20 across the four suggested thresholds. However, in the ECOLI network, average local clustering increased from 0.24 (low confidence) to 0.40 (high confidence), before decreasing down to 0.35 (highest confidence). In the YEAST network, the average local clustering remained stable around 0.27 between low and medium confidence, and then steadily increased to 0.36 at the highest confidence threshold before dropping off again (Figure 1B).

**Figure 1:**
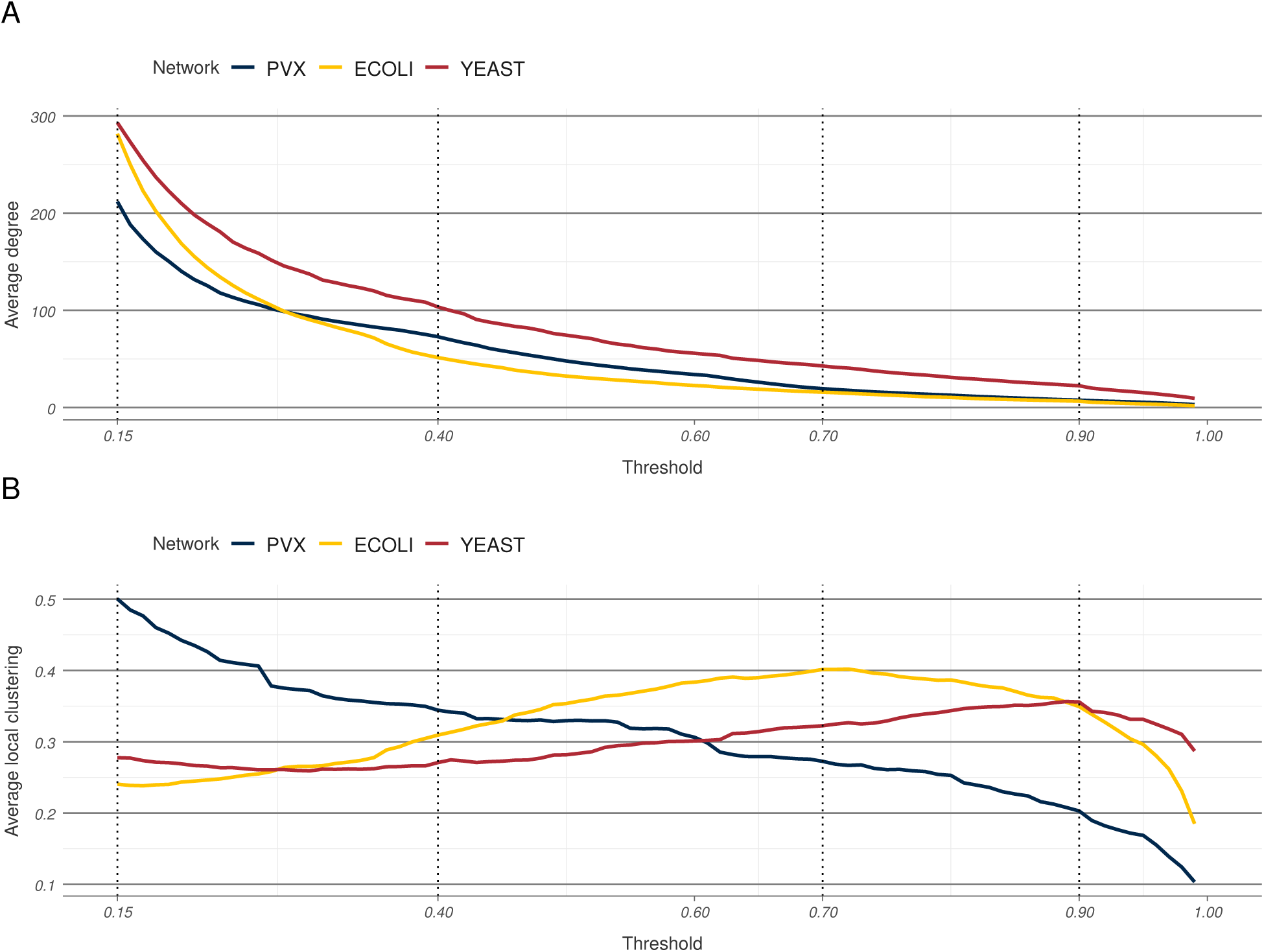
Thresholding effects in STRING networks. Average degree (A) and average local clustering coefficient (B) as functions of the threshold in the three STRING networks. The dotted vertical lines correspond to the four default STRING threshold values.

Figure 1 illustrates two aspects of scored PINs: firstly that node metrics can vary significantly in their raw values across thresholds, and secondly, that this variation is qualitatively different for different metrics. The incomplete coverage and experimental error of interaction detection techniques imply that it is most unlikely that any particular thresholded network describes perfectly all biologically relevant interactions and only them. Therefore, any robust, biologically informative PIN analysis pipeline should ideally show agreement in results obtained across at least a narrow range of different thresholds. In the context of using node metrics to identify key proteins in a network, such an agreement may translate to identifying the same set of highest ranking nodes.

We analysed 25 node metrics, of which 12 were node centralities, and 13 were global network summaries which we redefined as leave-one-out difference (LOUD) metrics. Four scored PINs were considered, spanning three organisms and two databases—the three PINs from STRING, illustrated in Figure 1 and the *S. cerevisiae* network obtained from HitPredict (HPRED). For each PIN, these metrics were calculated for all nodes in a set of 85 thresholded networks, obtained at equidistant thresholds from 0.15 (the lowest recorded) to 0.99 at 0.01 intervals. In addition we considered two synthetic scored networks—SYN-GNP based on a Bernoulli random graph, and SYN-PVX, based on a re-scored subset of the PVX network. The node rankings induced by a metric at each of the thresholded networks were used to assess the metric’s rank robustness.

### Rank continuity

PIN analysis often aims to identify key proteins in a particular biological process or context, for example by studying which nodes in the network attain high values across different node metrics. This problem relates to the node rankings induced by the metrics (*“Which are the nodes of highest degree?”*), rather than to exact metric values (*“What is the degree of these nodes?”*).

Exact metric values can be difficult to interpret and will vary both between PINs and with the PIN confidence score threshold. For example, ubiquitin (YLL039C) has degree between 262 and 4254 across different thresholds of the YEAST network (values obtained at score thresholds 0.99 and 0.15 respectively). These values are vastly different, and not easily interpretable or comparable outside the context of the particular thresholded networks they are obtained from. In contrast, the fact that ubiquitin is the single highest degree node across all thresholds for the scored YEAST network demonstrates its biological role more clearly.

We propose that for a node metric to be reliably indicative of the biological state described by a scored PIN, it should identify similar sets of key, i.e. highest ranking, proteins at a range of thresholds. In particular, rankings obtained at consecutive thresholds (e.g. at 0.40, the proposed medium confidence threshold in STRING, and at the slightly higher 0.41) should be in good agreement. Large differences could imply that (a) the metric is too influenced by pre-processing decisions to be informative, or (b) that the confidence score distribution is highly concentrated between these two thresholds and moving from one to the other significantly changes network topology.

For each analysed node metric, rank similarity was measured using Trajanovski’s *k*-similarity (see Methods) between each two consecutive thresholds, across all scored networks—the three STRING networks, the *S. cerevisiae* network from HitPredict, and two synthetic networks (Figures 2, S1-S4). In the analysed PINs, three different modes of behaviour were observed: some metrics exhibited consistently high similarity, some consistently low, and for others *k*-similarity steadily decreased with threshold increase (Figures 2A, S1 and S2).

**Figure 2:**
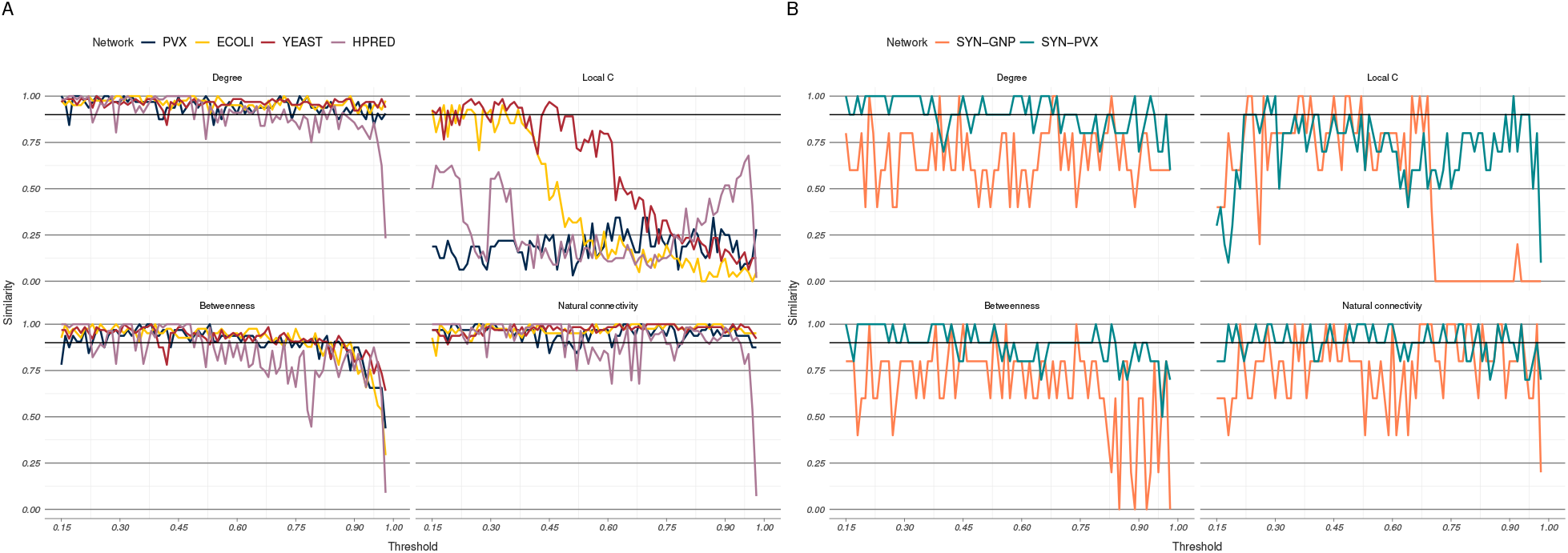
Metric rank similarity between consecutive thresholds. (A) In the four PINs, metrics were either consistently stable (e.g. degree and LOUD natural connectivity), consistently unstable (e.g. local clustering coefficient), or showed decreasing stability (e.g. betweenness). (B) The synthetic network based on a randomly rescored subset of the PVX network, SYN-PVX, and the network based on a Bernoulli random graph, SYN-GNP, exhibited different behaviour, with metrics showing the least similarity across thresholds in the SYN-GNP network.

We propose a rank continuity measure based on how often *k*-similarity between consecutive thresholds reaches 0.90 (details in Methods). Continuity was measured for our set of twenty-five metrics (Methods and SI for details) across the six scored networks (Tables S1 and S5). Between 7 and 16 metrics were found to have continuity measures over 0.90 for the medium-high confidence region in each of the scored PINs. The value of 0.90 was chosen since we believe robust metrics should produce nearly identical sets of high-ranking nodes across most pairs of consecutive thresholds. Seven metrics were found to have continuity measures over 0.90 in all four networks, and an additional four metrics had high continuity in three out of the four networks (all but the PVX network). Eleven of the twenty-five metrics—degree, redundancy, PageRank, harmonic closeness, LOUD natural connectivity, LOUD global clustering, LOUD average redundancy, and four of the ego-network based metrics—had an average score across the four PINs above 0.90. Nine metrics, including the commonly used local clustering coefficient and betweenness centrality, did not achieve a high continuity measure in any of the four PINs.

Spearman rank correlations of the continuity measures (Table S2) showed extremely good agreement between the three STRING networks (all correlations were above 0.95), and very good agreement between the STRING networks and the HPRED network (correlations between 0.89 and 0.92). These were higher than correlations between the STRING networks and either of the synthetic networks (between 0.30 and 0.68), suggesting that how edge scores are placed over the network (biased as opposed to random) plays an important role in metric rank continuity. Finally, continuity in the synthetic Bernoulli network, SYN-GNP, was considerably lower across all metrics (Figure 2B, Table S5). This implies that the metric continuity is sensitive to network structure—the reported continuity values will not necessarily hold for other types of networks (e.g. social, transport, etc.), where other node metrics may be more stable.

Our continuity analysis suggests that nearly half of the tested node metrics are robust to small threshold perturbation in PINs. However, incremental changes in the set of overlapping nodes between consecutive thresholds may result in a high continuity measure but low similarity between rankings at more distant thresholds. This can be undesirable, since often confidence scores are not readily interpretable and there may exist a wide range of permissible thresholds (e.g. anywhere between 0.15 and 0.90 in STRING).

### Rank identifiability

In order to assess the robustness of a node metric across a range of medium-high confidence score thresholds, overall ranks were calculated and compared to ranks induced at single thresholds. These ranks were designed to represent the relative position (i.e. importance) of each node across a range of medium-high confidence thresholds, and were calculated as rank averages across the threshold region. We define a rank identifiability measure, which quantifies the ability to recover the set of overall highest ranking nodes by considering any single threshold in the region (Methods). Our identifiability measure is based on an asymmetric version of *k*-similarity, which we introduce and call α-relaxed *k*-similarity (defined in Methods). Intuitively, a rank identifiability measure of 0.90 means that at least 90% of the overall highest ranking nodes are also ranked highly at any given threshold.

In each of the three STRING networks, 5 or 6 node metrics were found to have rank identifiability measures above 0.90 (Table S6). In the HPRED network, where the medium-high confidence region is shorter than in STRING, identifiability measures were higher and 16 metrics attained a score above 0.90. Of these, four metrics—redundancy, number of edges in the step-one ego network, LOUD natural connectivity, and LOUD average number of edges in the step-one ego network—had identifiability measures above 0.90 across all four PINs.

The similarities of rankings induced at thresholds outside the medium-high confidence region to the overall ranks were also calculated, although these similarities did not contribute to the rank identifiability scores. Since the overall ranks were calculated over the medium-high confidence region for each network, it is natural to expect α–relaxed *k*-similarities to be higher within the region than outside it (Figures 3, S5-S8). In the HPRED network, for example, some metrics showed high rank similarity for thresholds as high as 0.45. This indicates that the exact boundaries of the region do not necessarily heavily influence identifiability results.

**Figure 3:**
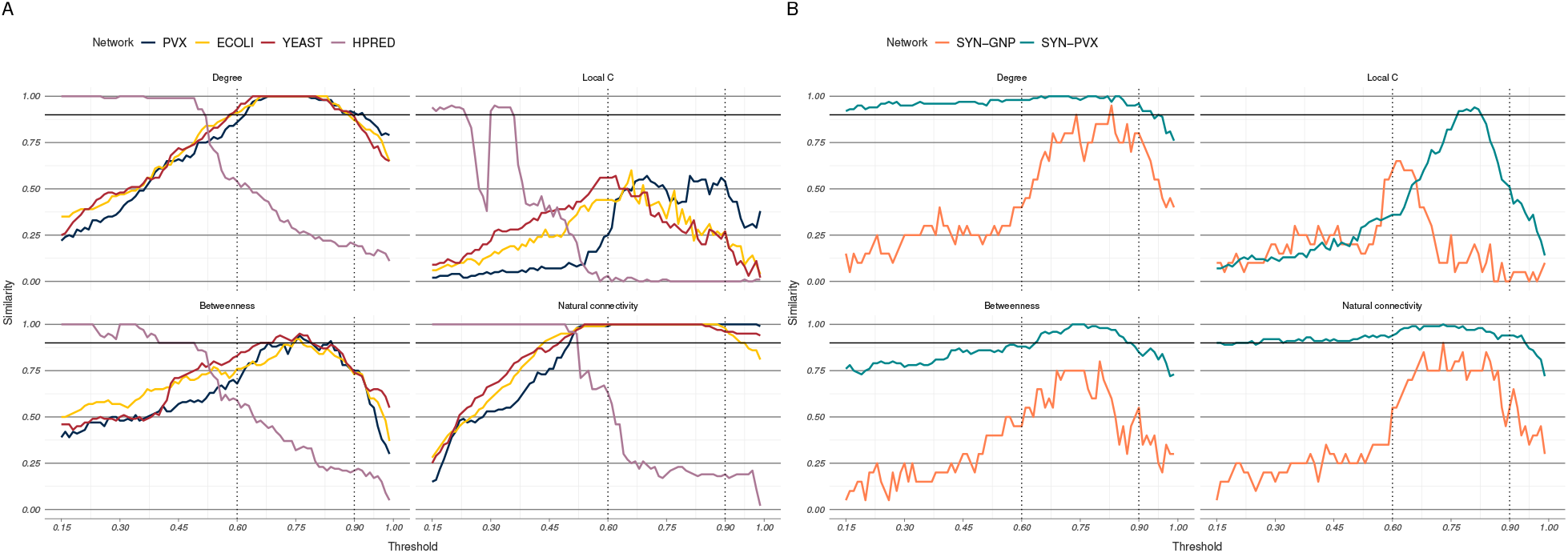
Relaxed similarity between overall and threshold ranks in the scored PINs. (A) The overall ranks have been calculated over the medium-high confidence regions—0.60 to 0.90 for the three STRING networks (black dotted lines) and 0.15 to 0.28 for the HitPredict network (pink dotted lines). (B) The STRING medium-high confidence region has also been used for the synthetic networks. The SYN-PVX network exhibits higher identifiability, likely because the randomisation of the edge scores has introdiced the same rate of change across the network.

Like rank continuity, rank identifiability is a context-dependent property of network metrics. The three STRING networks closely agreed in identifiability measures (Spearman correlations between metric identifiability scores were all 0.94 and above), and were more similar to measures obtained from analysing the HPRED network (correlations above 0.81) than any of the synthetic networks (Table S3).

While rank identifiability can be used to quantify rank conservation across many thresholds, it does not account for all types of rank variability. Intuitively, a metric which preserves the exact same ranking in the set of top *n* nodes at every threshold is more robust than one in which the top node set is preserved, but re-ranked. However, since our rank continuity and identifiability measures are both based on set overlap only, in both the preserved and the re-ranked case the metrics would achieve a perfect score of 1. To take this difference into account we introduce a measure for rank instability.

### Rank instability

A different way of assessing how well top-ranking nodes (or nodes in general) preserve their ranks across different thresholds is to calculate the ranges of ranks they attain. A robust metric should result in relatively narrow rank ranges. In particular, overall top ranking nodes should have relatively narrow rank ranges.

In order to quantify this behaviour we define rank instability as the scaled average rank range of the overall top 1% ranking nodes (details in Methods). Unlike the rank continuity and identifiability measures, where values close to one represent robustness, instability values close to zero correspond to narrower rank ranges, and therefore more consistent node metric behaviour. The instability measure was lower in the HPRED network, where the medium-high confidence interval is shorter, and similar across the three STRING PINs (Figure 4A and Table S7). Only four metrics had instability measures below 0.01 in all PINs—number of edges in the step-1 ego network, LOUD natural connectivity, LOUD average redundancy and LOUD average number of edges in the step-1 ego network.

**Figure 4:**
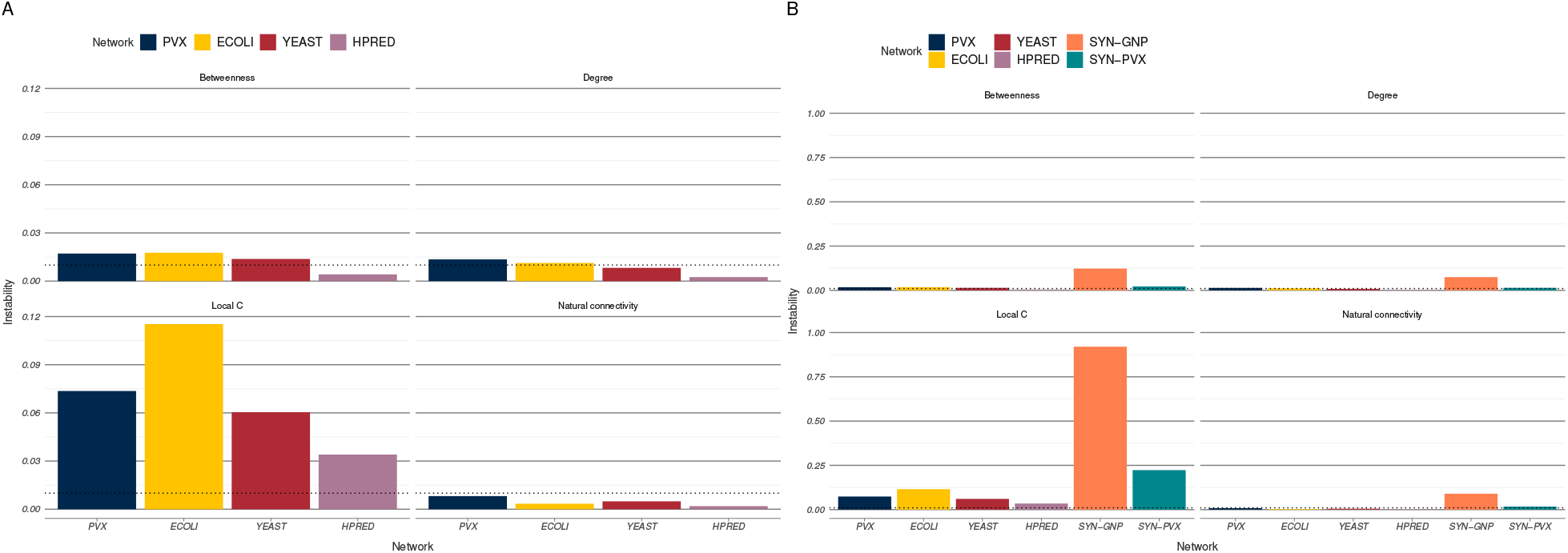
Rank instability of metrics in the scored networks. (A) Rank instability in the four PINs. The dotted lines correspond to 1%. Instability measures in the HPRED network were generally narrower. (B) Rank instability in the synthetic networks. Instability measures in PINs were generally lower and have been plotted for comparison. Note the different scales between plots in A and in B.

Rank instability measures in the synthetic networks were generally higher than in the scored PINs (Figure 4B and Table S7). Spearman correlations of metric rank instability were higher across the scored PINs than they were between PINs and either of the synthetic networks (Table S4), suggesting once again that PINs exhibit context-specific behaviour.

Overall, the three measures for rank robustness of node metrics described here—rank continuity, identifiability, and instability—agree across the four studied protein interaction networks and identify four node metrics to be robust to thresholding: number of edges in the step-1 ego network, LOUD natural connectivity, LOUD average redundancy and LOUD average number of edges in the step-1 ego network. Measures of rank continuity, identifiability, and instability for all 25 analysed metrics averaged over the four PINs can be found in Table S1.

## Discussion

Protein interaction network analysis typically starts with network construction—some interaction data of interest is obtained, either experimentally or from publicly available databases, and is then pre-processed before being used to create a network. Different types of protein interactions, experimental systems for detecting them, and data clean-up choices mean that multiple networks can be built to represent the same underlying biological process. Since typically only one of these will be analysed, it is not immediately clear how much these network models of cellular biology and any conclusions drawn from them might differ. In this paper we have demonstrated that varying the confidence score threshold for interactions can result in structurally different networks, both in terms of density and in terms of properties like average local clustering coefficient. We propose that if PIN analysis aims to provide reliable, reproducible biological insight, it should show some agreement across alternative network models.

In this paper, we have focused on one frequent goal of PIN analysis—identifying key proteins (nodes) in a particular piece of biological architecture (network). This can be done by calculating node metrics, such as degree or betweenness centrality, and then identifying the set of nodes which rank highest based on metric performance. There are many different metrics one can use in this context, and it is not always clear which are the most suitable. We argue that one desirable feature a suitable metric should posses is rank robustness, or the ability to identify the same or at least largely similar sets of top ranking proteins when the network construction process is altered. Here we have considered robustness to variation in the confidence score threshold required to include an interaction in the network.

Networks constructed at lower thresholds are denser and potentially include more spurious interactions than networks constructed at higher thresholds. Increasing the stringency of data quality, however, may result in potentially important but under-studied interactions to be omitted. Different scoring procedures across databases, and even within different versions of the same database (Figure S9), make optimal threshold choice difficult. Therefore, a level of rank robustness across different thresholds is desirable for node metrics. We have proposed three measures with which to assess such robustness—rank continuity, identifiability, and instability. The relevance of each measure will depend on the research question at hand and on the reliability of the confidence scoring procedure. Rank continuity captures the effect of small threshold perturbations. This measure may be of particular interest when a narrow band of permissible thresholds has been identified. Rank identifiability compares threshold ranks at given thresholds against overall ranks. It may be particularly informative when there are no known optimal threshold values. Finally, rank instability assesses the variation in threshold ranks for the overall top 1% of nodes. It is our most stringent measure, requiring not only that highly ranked proteins remain highly ranked but also that they retain their ordering across thresholds. It should be used when absolute rank is important.

We calculated the rank continuity, identifiability, and instability of 25 node metrics in each of six scored networks, four of which were PINs and two of which were synthetically generated. Our rank continuity measure, which is based on a rank similarity measure originally proposed by [42], quantifies the agreement between node rankings obtained at consecutive thresholds. If networks obtained at two close thresholds, say 0.50 and 0.51, yield considerably different rankings, this may suggest that threshold choice plays an overwhelmingly important role in network construction, and may obscure any underlying biological signal that could otherwise be detected. Conversely, high continuity measures correspond to rankings which are unlikely to significantly change with small threshold perturbations.

Confidence scores are not always readily interpretable, which might make threshold choice more difficult. Therefore, agreement over a wider threshold region might also be desirable. Small differences in consecutive thresholds might be responsible for large discrepancies between more distant thresholds (say 0.50 and 0.70), while still preserving a high rank continuity measure. In order to take this effect into account, we introduce rank identifiability to measure the agreement between different threshold rankings and a single overall ranking.

Finally, our rank instability measure provides an alternative way of analysing rank robustness which is not based on rank overlap but instead focuses on the different ranks a particular node attains at different thresholds. High instability corresponds to the overall top ranking nodes attaining a wide range of individual threshold ranks.

Our analysis identified four node metrics—number of edges in the step-one ego network, LOUD average redundancy, LOUD average number of edges in the step-one ego network, and LOUD natural connectivity— which induce robust ranks across all four analysed PINs. More commonly used metrics such as degree, local clustering coefficient, betweenness, and closeness did not perform as well. For example, when comparing the top 100 nodes obtained at the start and the end of the medium-high confidence region for the YEAST network (thresholds set at 0.60 and 0.90 respectively), node sets obtained using LOUD natural connectivity showed a three quarter overlap (75 out of 100). In contrast, the overlap of the top ranking sets identified by betweenness was less than half (41 out of 100), and the overlap between sets identified using local clustering coefficient was less than 10% (9 out of 100).

Spearman rank correlations between robustness measures across the four different PINs were consistently high (0.89 and above). The analysed PINs varied in organism, database, confidence scoring methodology, and even interaction type (functional associations in STRING, and physical interactions only in HitPredict). This implies that the presented robustness results may be readily applicable to other PINs. In contrast, the lower correlations observed when scored PINs were compared with the two synthetic networks indicate that rank robustness is context-specific.

## Conclusions

Protein interaction data can be obtained using a range of techniques and is subject to different types of experimental error. The uncertainty associated with interaction detection can be quantified by confidence scores. Applying a threshold to these scores provides researchers with a degree of control over the false positives and false negatives present in protein interaction networks. Nevertheless, we argue that if PIN analysis aims to capture biological insight, it should show a level of agreement across networks obtained at different thresholds.

We present a methodology for assessing metric rank robustness. We demonstrate that network topology will generally vary with threshold choice, but that certain node metrics remain relatively stable. This dependence on threshold choice will affect metric interpretability in the context of bioinformatics research.

## Materials and Methods

### Protein interaction and synthetic networks

In order to assess the rank robustness of different network metrics, four scored protein interaction networks were used. The networks ranged across two databases and three organisms. A confidence score quantifying the reliability of available interaction evidence was available for each detected edge across all four networks.

Three organism networks—*P. vivax* (retrieved March 2018), *E. coli*, and *S. cerevisiae* (both retrieved Feb 2018)—were obtained from STRING [28]. STRING contains both physical and functional association data, collected across a range of experimental and *in silico* interaction detection techniques. The organisms were chosen because they are model organisms with higher-than-average coverage of protein-protein interaction screens, while also having relatively short genomes, thus reducing the computational cost of our analysis.

In order to allow for a comparison between databases, the interaction network for *S. cerevisiae* was also downloaded from HitPredict [29]. Unlike STRING, HitPredict is a curated database containing only high-confidence physical interactions.

Filters were applied to remove duplicate interaction records, self-interactions, and interactions to proteins of other organisms. Only combined interaction scores were considered in all four cases, ignoring any available sub-scores. Confidence score distributions for the four networks can be seen in Figure 5.

**Figure 5:**
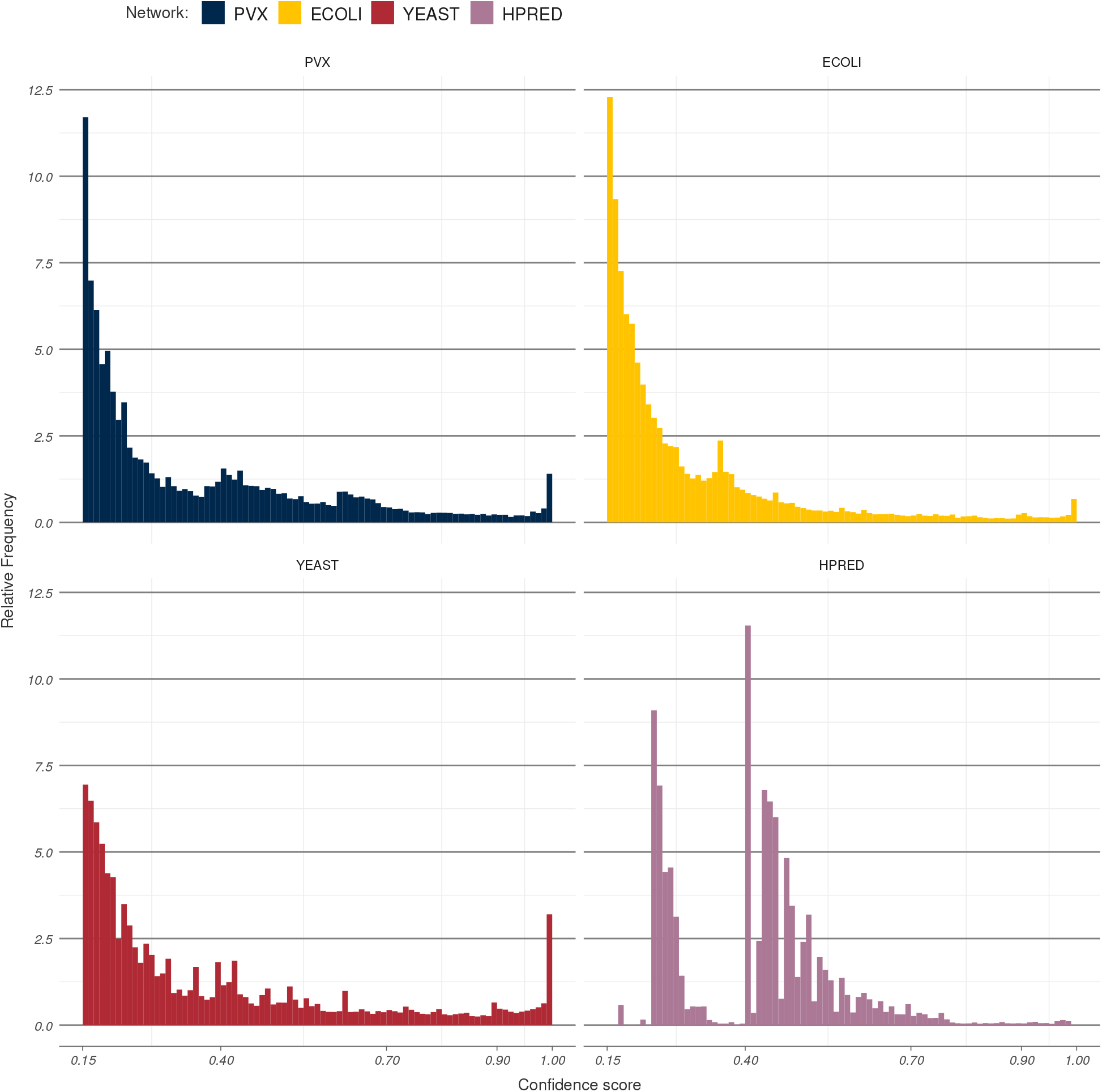
Confidence score distributions in each of the four studied PINs. Bin width in all four cases has been set to 0.01. Scores from the HitPredict network (bottom right) follow a different distribution and cannot necessarily be interpreted in the same way as STRING scores.

Two synthetic scored networks were also analysed. One was generated using a Bernoulli random graph model on *N* = 500 nodes and with edges sampled independently at random with probability *p* = 0.06. The value of *p* was chosen to be close to network density in the STRING PINs before thresholding. Edge scores were sampled with replacement from the *P. vivax* confidence score distribution. The second network was an induced random subgraph of the *P. vivax* network on *N* = 1000 nodes. The available edge scores for the subgraph were randomly rearranged and placed over the fixed edges. This resulted in a network which is PIN-like in topology, but which contains no local dependency between edge scores. A summary of all six analysed networks can be found in Table 1.

**Table 1:** Summary statistics for the six analysed networks. The left-most column corresponds to the names the networks are referred as later in the text. The number of edges and edge density refer to the all scored edges before any threshold is applied to the network.

| Name | Network | Number of nodes | Number of edges | Edge density |
| --- | --- | --- | --- | --- |
| PVX | *P. vivax*, STRING | 3255 | 344691 | ~0.065 |
| ECOLI | *E. coli*, STRING | 4144 | 583440 | ~0.068 |
| YEAST | *S. cerevisiae*, STRING | 6418 | 939998 | ~0.046 |
| HPRED | *S. cerevisiae*, HitPredict | 5673 | 113001 | ~0.007 |
| SYN-GNP | Synthetic, Bernoulli | 500 | 7459 | ~0.060 |
| SYN-PVX | Synthetic, randomised *P. vivax* | 1000 | 30516 | ~0.061 |

### Thresholding

Applying a threshold *θ* to a scored PIN means discarding all edges in the network with scores strictly lower than *θ* and creating a simple, unweighted network from the remaining edges. The node set is not altered. Figure 6 gives a schematic of how thresholds are applied.

**Figure 6:**
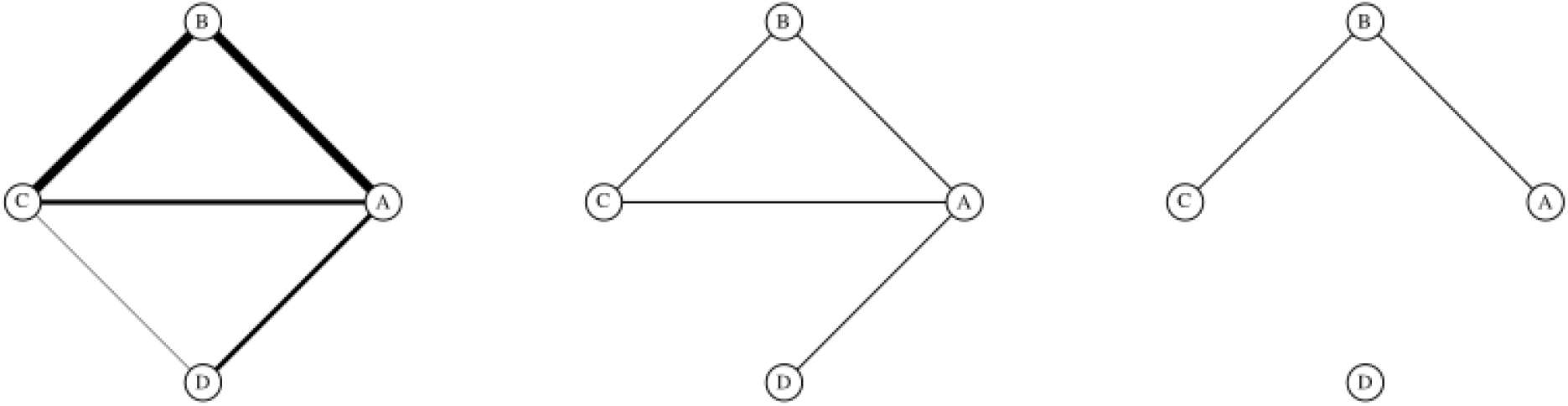
Thresholding scored networks. A scored network, with edge widths corresponding to confidence scores (left). At a low threshold, only the lowest scoring edge CD is removed (middle). At a higher threshold, only the highest scoring edges AB and BC remain in the network (right). Edge scores are otherwise ignored in the thresholded networks.

All reported confidence scores in the analysed PINs were between 0.15 and 1.00. Thresholds were applied from 0.15 to 0.99 inclusive at 0.01 intervals, resulting in a set of 85 distinct thresholded networks for each of the scored PINs. The same node set was preserved across all 85 thresholded networks, even when thresholding resulted in isolating nodes from the rest of the network. In order to minimise the effect of extreme thresholding, both the full threshold region and a truncated, medium-high confidence region were analysed for each network.

The majority of interactions in STRING are re-scored across different database releases (Figure S9) which indicates that the scores themselves should be treated with some error tolerance. In order to take this into account while retaining score interpretability, a wide medium-high confidence region was set between 0.60 and 0.90. Medium-high confidence scores in STRING occur at similar, low frequencies across organisms (Figure 5). HitPredict generally contains higher quality data and employs a different, more stringent scoring procedure—while interaction scoring 0.40 in STRING would be considered medium confidence, the same score in HitPredict would indicate high confidence. The truncated medium-high confidence region for the HPRED network was therefore set between 0.15 and 0.28, since HitPredict scores above 0.28 are considered to be high-confidence [29].

### Metric extraction and ranking

The rank robustness of twenty-five node metrics was studied. These included twelve node centralities, and thirteen leave-one-out difference (LOUD) global network summaries.

Commonly used metrics, such as degree, local clustering coefficient, betweenness and closeness centralities were included in the node metric set. In addition, metrics based on the size and density of the step-one and step-two ego networks for each node were calculated. The step-one ego network for a node *v* was formed by taking the subgraph induced by the *v* and its immediate neighbours (graph distance 1); in addition, the step-two ego-network also included nodes within graph distance 2 from *v*.

The LOUD metrics were used as a way to assess the effect of perturbing the network by isolating each node in turn. These included, wherever applicable, local metrics averaged over the entire network (e.g. average local clustering coefficient), as well as metrics which are by definition global (e.g. global clustering coefficient). The set also included natural connectivity, a spectral metric designed to measure the overall robustness of a network [44]. Due to the associated computational costs, LOUD metrics were only calculated for nodes with degree at least two. Metric values for leaves and isolated nodes were set to NA (not available).

The nodes in each thresholded network were ranked by each of the metrics, with high ranks corresponding to high metric values. Node ranks for nodes for which LOUD metrics were not evaluated were set to first, i.e. smallest. Ties were resolved at random, independently across different metrics and different thresholds. A full list of node metrics, as well as details on their computation, can be found in the SI.

### Evaluation of rank robustness

As the threshold applied to the scored networks increases, the networks become lower in edge density and node metric values will be affected—for example, node degrees will decrease. In order to assess the robustness of each metric to changes in threshold, the node rankings induced by the metric at different thresholds were compared instead of the calculated values.

Rank similarity is typically measured by a rank correlation coefficient such as Spearman or Kendall. These coefficients are used to compare whole rank vectors. In the context of bioinformatics applications, node metrics are often used to identify the key actors in a particular process, and therefore it is natural to focus on the highest ranking nodes only. In order to do this, robustness analysis was based on the rank similarity measure *M_k_* proposed by [42] as follows.

A ranking *A* is a vector *A* = {*A_v_* : *v* ∈ {1, …, *N*}} of ranks assigned to the network nodes *v* ∈ {1, …, *N*}, e.g. by considering their degree at a particular threshold. The *k*-similarity of two rankings *A*_1_ and *A*_2_ is the overlap between their 100*k*% highest ranking nodes, 
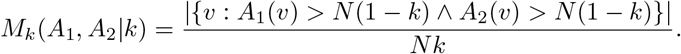

This measure of rank similarity is symmetric, and is therefore useful for cases where both rankings carry the same meaning, e.g. when they are obtained at consecutive thresholds. However, it is too restrictive when the rankings being compared are interpreted differently. To account for this, we introduce an *α*-relaxed asymmetric version of *k*-similarity. The *α*-relaxed *k*-similarity of a ranking *A* to some ranking *B* is the proportion of the top 100*k*% highest ranking nodes in *B* which are also within the set of 100(*k* + *α*)% highest ranking nodes in *A*: 
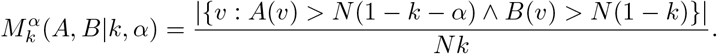

Our relaxed *k*-similarity allows for more user control when the rankings compared are not interpreted in the same way, and need therefore not be treated in the same way. For example, if *A* is obtained from a single threshold, and *B* is some overall ranking, relaxed *k*-similarity may be used to identify whether the top 10 nodes overall, i.e. in *B*, are among the top 20 for the particular observed threshold, i.e. in *A*.

### Rank continuity

We introduce *rank continuity* of each metric in each network, i.e. the similarity between rankings induced at consecutive thresholds. In all cases a set of values for the proportion of nodes k considered to be the top ranking were used, ranging from 0.001 to 0.05 at 0.001 intervals. An overall continuity measure was calculated based on how often the observed similarity was high (0.90 or above).

Suppose that a metric *f* induces node rankings *A_i_*, *A_i_*_+0.01_, …, *A_j_* at each threshold within the medium-high confidence region. Then we define the rank continuity of *f* as the fraction of cases where the *k*-similarity between consecutive *A_i_* and *A_i_*_+0.01_ is over 0.90: 
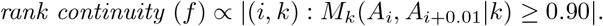

Since a range of different values of k up to 0.05 was considered in calculating a single continuity measure, higher ranking nodes contribute more to the overall rank continuity of a metric.

### Rank identifiability

Further, for each metric an overall ranking was calculated for the truncated threshold region. The overall ranks for all nodes were calculated by first averaging over node ranks at all relevant thresholds, and then ranking the resulting values. Ties were resolved at random.

For our definition of rank identifiability, for each metric the *α*-relaxed *k*-similarity between overall ranks *B* and threshold ranks *A_i_* for each threshold *i* in the medium-high confidence region was calculated. The rank identifiability score for each metric *f* was defined as the minimum observed relaxed similarity between overall and threshold ranks.

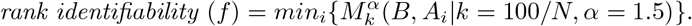

In all cases apart from the Bernoulli network, which had the smallest number of nodes, the ability to recover the top *n* = 100 nodes overall (i.e. *k* = 100*/N*) among the top 150 (i.e. *α* = 1.5) at any threshold was tested. In the Bernoulli network, *n* was lowered to *n* = 20, and the parameter *α* remained fixed at *α* = 1.5.

### Rank instability

Another way to assess rank robustness is through the *instability* of node ranks attained by different thresholds.

Rank ranges attained by the top 1% overall top ranking nodes over the medium-high confidence region were calculated for all scored networks. Ranges were scaled to the number of nodes in each network. Finally, the average of the scaled rank ranges was calculated for each metric, giving the instability measure for the metric.

A worked example of metric extraction, ranking, and robustness analysis can be found in the SI.

### Availability of data and materials

The datasets generated and analysed during the current study and accompanying software are available on the Oxford Protein Informatics Group website http://opig.stats.ox.ac.uk/resources.

## Author’s contributions

LVB, GR and CMD designed the study. LVB developed the rank robustness measures, carried out data analysis, and wrote the manuscript. GR and CMD supervised the work and edited the manuscript. AVW and JW contributed to node metric choice and data acquisition, and critically reviewed the manuscript. All authors reviewed and approved the final version of the manuscript.

## Acknowledgements

We thank Javier Pardo Díaz for critical reading of an earlier draft of this manuscript.

