## Supplementary information for "Measuring rank robustness in scored protein interaction networks"

### Measuring rank robustness in scored protein interaction networks: supplementary information

Lyuba V. Bozhilova[1], Alan V. Whitmore[2], Jonny Wray[2], Gesine Reinert[1], Charlotte M. Deane[1*]
**1** Department of Statistics, University of Oxford, Oxford, OX1 3LB, United Kingdom
**2** e-Therapeutics Plc, 17 Fenlock Rd, Long Hanborough, OX29 8LN, United Kingdom
*

### Contents

**1 Results across all four networks** **2**

**2 Rank continuity: additional results** **4**

**3 Rank identifiability: additional results** **9**

**4 Rank instability: additional results** **14**

**5 STRING network v.9 and v.10** **14**

**6 Metric extraction** **16**

### 1 Results across all four networks

| Metric | Continuity | Identifiability | Instability |
|---|---|---|---|
| Degree | **0.96** | 0.90 | 0.01 |
| Local clustering | 0.21 | 0.28 | 0.07 |
| Redundancy | 0.92 | **0.97** | 0.01 |
| **Ego1: edges** | **0.96** | **0.96** | **<0.005** |
| Ego2: nodes | 0.84 | 0.79 | 0.02 |
| Ego2-Ego1: nodes | 0.71 | 0.74 | 0.02 |
| Ego1$^2$-Ego2: nodes | **0.96** | 0.90 | 0.01 |
| Ego1/Ego2: nodes | 0.17 | 0.12 | 0.23 |
| PageRank | 0.94 | 0.89 | 0.01 |
| Closeness | 0.86 | 0.83 | 0.01 |
| Harmonic centrality | 0.93 | 0.84 | 0.01 |
| Betweenness | 0.64 | 0.77 | 0.01 |
| **LOUD Natural connectivity** | **0.96** | **0.98** | **<0.005** |
| LOUD Avg. local clustering | 0.31 | 0.45 | 0.07 |
| LOUD Global clustering | 0.94 | 0.94 | 0.01 |
| **LOUD Avg. redundancy** | **0.96** | **0.95** | **<0.005** |
| **LOUD Avg. Ego1: edges** | **0.97** | **0.96** | **<0.005** |
| LOUD Avg. Ego2: nodes | 0.87 | 0.82 | 0.01 |
| LOUD Avg. Ego2-Ego1: nodes | 0.79 | 0.78 | 0.01 |
| LOUD Avg. Ego1$^2$-Ego2: nodes | **0.96** | 0.91 | 0.01 |
| LOUD Avg. Ego1/Ego2: nodes | 0.50 | 0.44 | 0.05 |
| LOUD No connected node pairs | 0.05 | 0.34 | 0.35 |
| LOUD Avg. shortest path | 0.32 | 0.28 | 0.09 |
| LOUD Avg. betweenness | 0.31 | 0.21 | 0.11 |
| LOUD Avg. closeness | 0.32 | 0.66 | 0.03 |

Table S1: **Continuity, identifiability and instability measures for all metrics, averaged across the four PINs.** Metrics for which average continuity and identifiability were above 0.95 and average instability was below 0.005 are highlighted.

| | PVX | ECOLI | YEAST | HPRED | SYN-GNP | SYN-PVX |
|---|---|---|---|---|---|---|
| PVX | 1.00 | 0.95 | 0.96 | 0.92 | 0.46 | 0.48 |
| ECOLI | 0.95 | 1.00 | 0.96 | 0.92 | 0.58 | 0.54 |
| YEAST | 0.96 | 0.96 | 1.00 | 0.89 | 0.56 | 0.57 |
| HPRED | 0.92 | 0.92 | 0.89 | 1.00 | 0.42 | 0.30 |
| SYN-GNP | 0.46 | 0.58 | 0.56 | 0.42 | 1.00 | 0.79 |
| SYN-PVX | 0.48 | 0.54 | 0.57 | 0.30 | 0.79 | 1.00 |

Table S2: **Spearman correlations between rank continuity measures across the different networks.** High correlations between the protein interaction networks, and lower correlations with the synthetic networks suggest that metric rank continuity depends both on network topology and on score placement.

| | PVX | ECOLI | YEAST | HPRED | SYN-GNP | SYN-PVX |
|---|---|---|---|---|---|---|
| PVX | 1.00 | 0.96 | 0.94 | 0.85 | 0.42 | 0.62 |
| ECOLI | 0.96 | 1.00 | 0.97 | 0.84 | 0.34 | 0.54 |
| YEAST | 0.94 | 0.97 | 1.00 | 0.81 | 0.37 | 0.57 |
| HPRED | 0.85 | 0.84 | 0.81 | 1.00 | 0.46 | 0.60 |
| SYN-GNP | 0.42 | 0.34 | 0.37 | 0.46 | 1.00 | 0.78 |
| SYN-PVX | 0.62 | 0.54 | 0.57 | 0.60 | 0.78 | 1.00 |

Table S3: **Spearman correlations between rank identifiability measures across the different networks.** Similar to rank continuity (Table EV1), correlations are highest between the scored PINs. The SYN-PVX network correlates to the scored PINs better than the SYN-GNP network does.

| | PVX | ECOLI | YEAST | HPRED | SYN-GNP | SYN-PVX |
|---|---|---|---|---|---|---|
| PVX | 1.00 | 0.95 | 0.94 | 0.91 | 0.40 | 0.64 |
| ECOLI | 0.95 | 1.00 | 0.97 | 0.92 | 0.37 | 0.62 |
| YEAST | 0.94 | 0.97 | 1.00 | 0.92 | 0.39 | 0.61 |
| HPRED | 0.91 | 0.92 | 0.92 | 1.00 | 0.26 | 0.61 |
| SYN-GNP | 0.40 | 0.37 | 0.39 | 0.26 | 1.00 | 0.83 |
| SYN-PVX | 0.64 | 0.62 | 0.66 | 0.61 | 0.83 | 1.00 |

Table S4: **Spearman correlations between rank instability measures across the different networks.** Correlations are again highest between the scored PINs. Like rank continuity and rank identifiability, (Tables EV2 and EV3), rank instability measures are more highly correlated between the two synthetic networks than they are between the SYN-GNP network and the scored PINs.

#### 2 Rank continuity: additional results

| Metric | PVX | ECOLI | YEAST | HPRED | SYN-GNP | SYN-PVX |
|---|---|---|---|---|---|---|
| LOUD Avg. betweenness | 0.18 | 0.02 | 0.40 | 0.62 | 0.09 | 0.26 |
| LOUD Avg. closeness | 0.15 | 0.14 | 0.32 | 0.66 | 0.10 | 0.36 |
| LOUD Avg. Ego1: edges | **0.92** | **0.97** | **0.99** | **0.98** | 0.12 | 0.54 |
| LOUD Avg. local clustering | 0.16 | 0.05 | 0.34 | 0.69 | 0.09 | 0.22 |
| LOUD Avg. Ego2-Ego1: nodes | 0.49 | **0.91** | **0.98** | 0.78 | 0.18 | 0.62 |
| LOUD Avg. Ego1/Ego2: nodes | 0.39 | 0.34 | 0.65 | 0.63 | 0.12 | 0.25 |
| LOUD Avg. $\text{Ego1}^2$-Ego2: nodes | **0.91** | **0.96** | **0.99** | **0.98** | 0.26 | 0.65 |
| LOUD Avg. Ego2: nodes | 0.74 | **0.94** | **0.97** | 0.82 | 0.26 | 0.63 |
| LOUD Avg. shortest path | 0.22 | 0.02 | 0.43 | 0.63 | 0.10 | 0.34 |
| LOUD Avg. redundancy | **0.92** | **0.96** | **0.99** | **0.99** | 0.07 | 0.15 |
| Betweenness | 0.54 | 0.55 | 0.69 | 0.77 | 0.09 | 0.54 |
| Closeness | 0.70 | 0.85 | **0.96** | **0.93** | 0.12 | 0.49 |
| Harmonic centrality | 0.84 | **0.93** | **0.98** | **0.97** | 0.11 | 0.57 |
| Degree | **0.91** | **0.96** | **0.99** | **0.97** | 0.10 | 0.59 |
| Ego1: edges | **0.92** | **0.96** | **0.99** | **0.99** | 0.10 | 0.52 |
| LOUD Global clustering | 0.83 | 0.95 | **0.98** | **0.99** | 0.13 | 0.12 |
| Local clustering | 0.14 | ∼0 | 0.25 | 0.43 | 0.07 | 0.17 |
| Ego2-Ego1: nodes | 0.38 | 0.89 | **0.93** | 0.63 | 0.12 | 0.54 |
| Ego1/Ego2: nodes | ∼0 | 0.02 | 0.04 | 0.62 | 0.04 | ∼0 |
| $\text{Ego1}^2$-Ego2: nodes | **0.91** | **0.96** | **0.99** | **0.97** | 0.31 | 0.69 |
| Ego2: nodes | 0.64 | **0.90** | **0.95** | 0.85 | 0.16 | 0.56 |
| LOUD Natural connectivity | **0.92** | **0.96** | **0.99** | **0.99** | 0.25 | 0.52 |
| LOUD No connected node pairs | 0.05 | 0.01 | 0.04 | 0.08 | 0.06 | 0.08 |
| PageRank | 0.87 | **0.95** | **0.99** | **0.95** | 0.31 | 0.67 |
| Redundancy | 0.83 | **0.94** | **0.95** | **0.98** | 0.08 | 0.11 |

Table S5: **Rank continuity for the 25 metrics across each of the six scored networks.** Values above 0.90 have been highlighted. The PINs show good general agreement, and the SYN-GNP network consistency exhibits lower rank continuity.

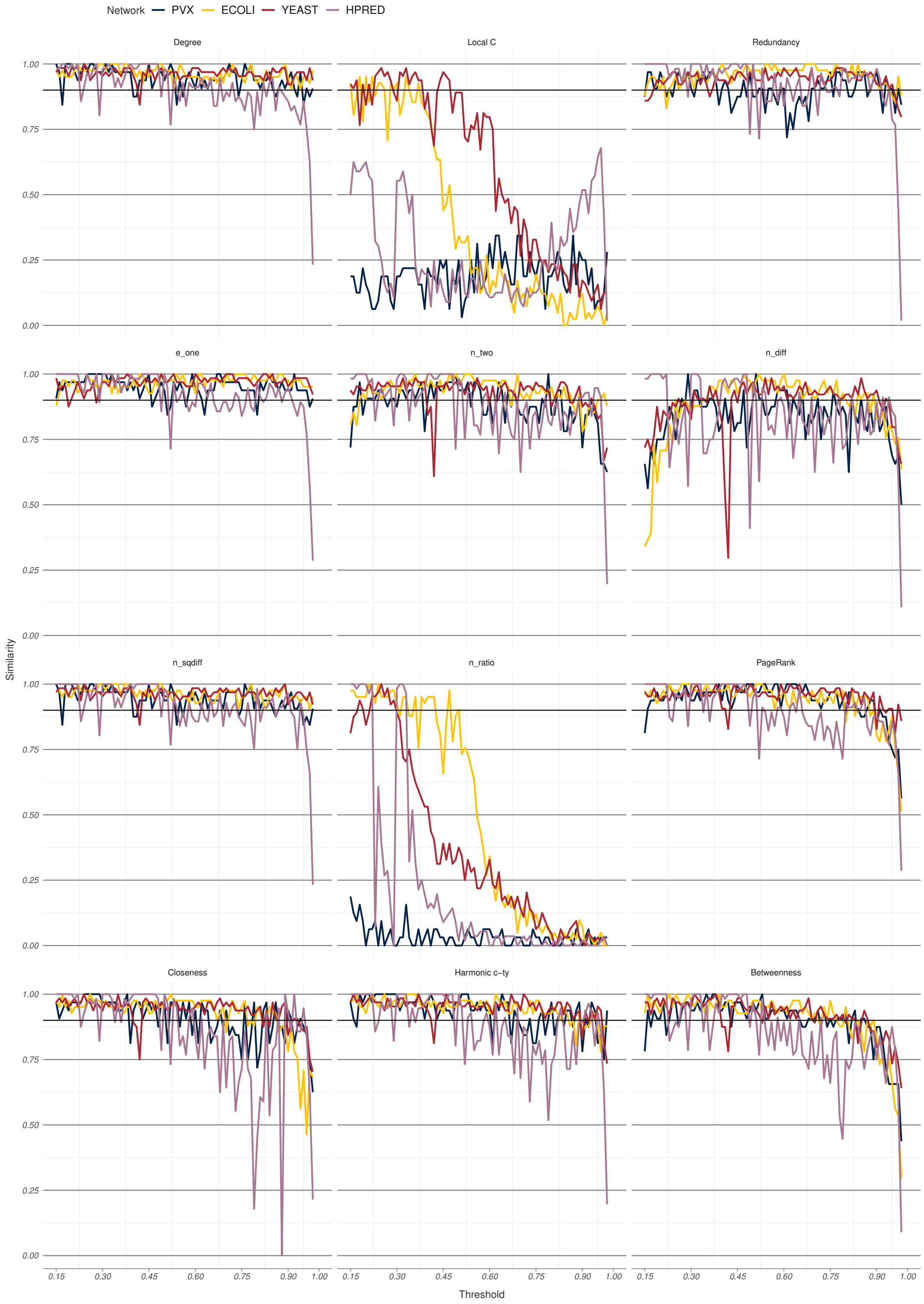

Figure S1: **Standard metric rank similarity between consecutive thresholds for the four PINs.** PINs across different species and databases showed generally good agreement. Local clustering coefficient (Local C) and the ratio between step-1 and step-2 ego networks (n_ratio) perform noticeably worse than other metrics.

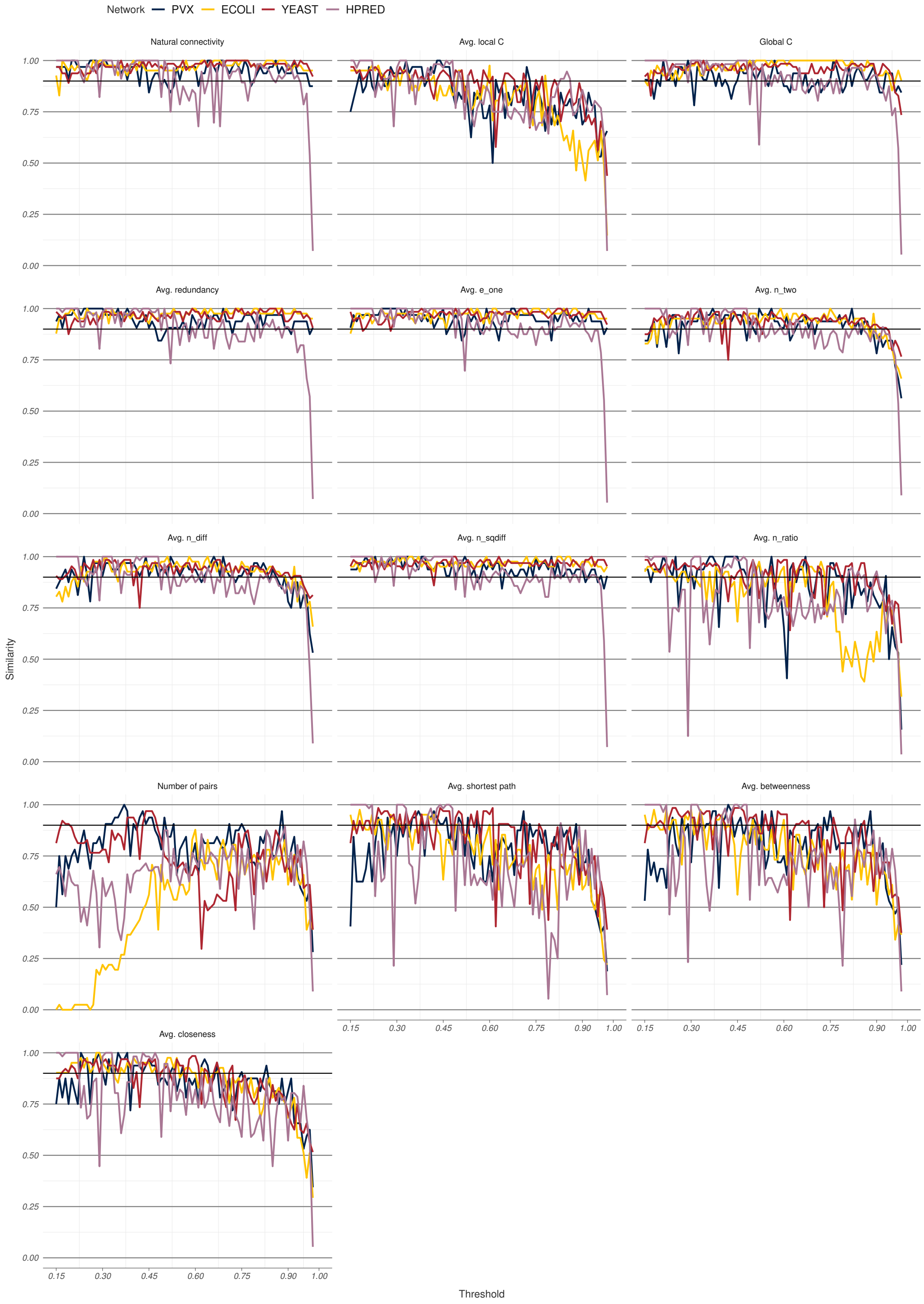

Figure S2: **LOUD metric rank similarity between consecutive thresholds for the four PINs.** Protein interaction networks across different species and databases showed generally good agreement. Average local clustering coefficient, average shortest path, average betweenness, and average closeness all exhibit a similar pattern of decreasing k-similarity as the threshold increases.

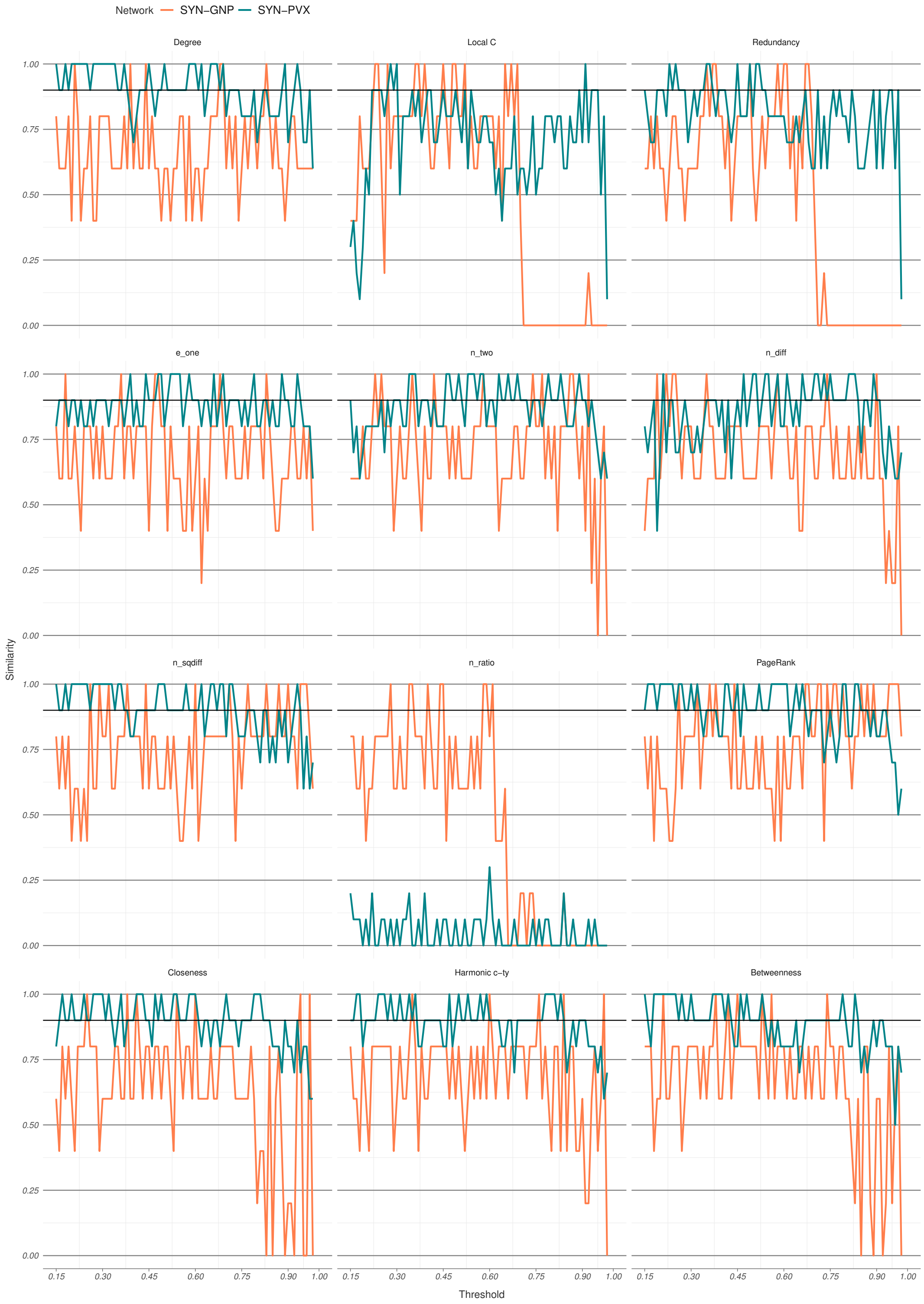

Figure S3: **Standard metric rank similarity between consecutive thresholds for the synthetic networks.** The Bernoulli synthetic network, SYN-GNP, exhibits consistently lower similarity across all node metrics aside from the ratio between the step-1 and step-2 ego networks (n_ratio).

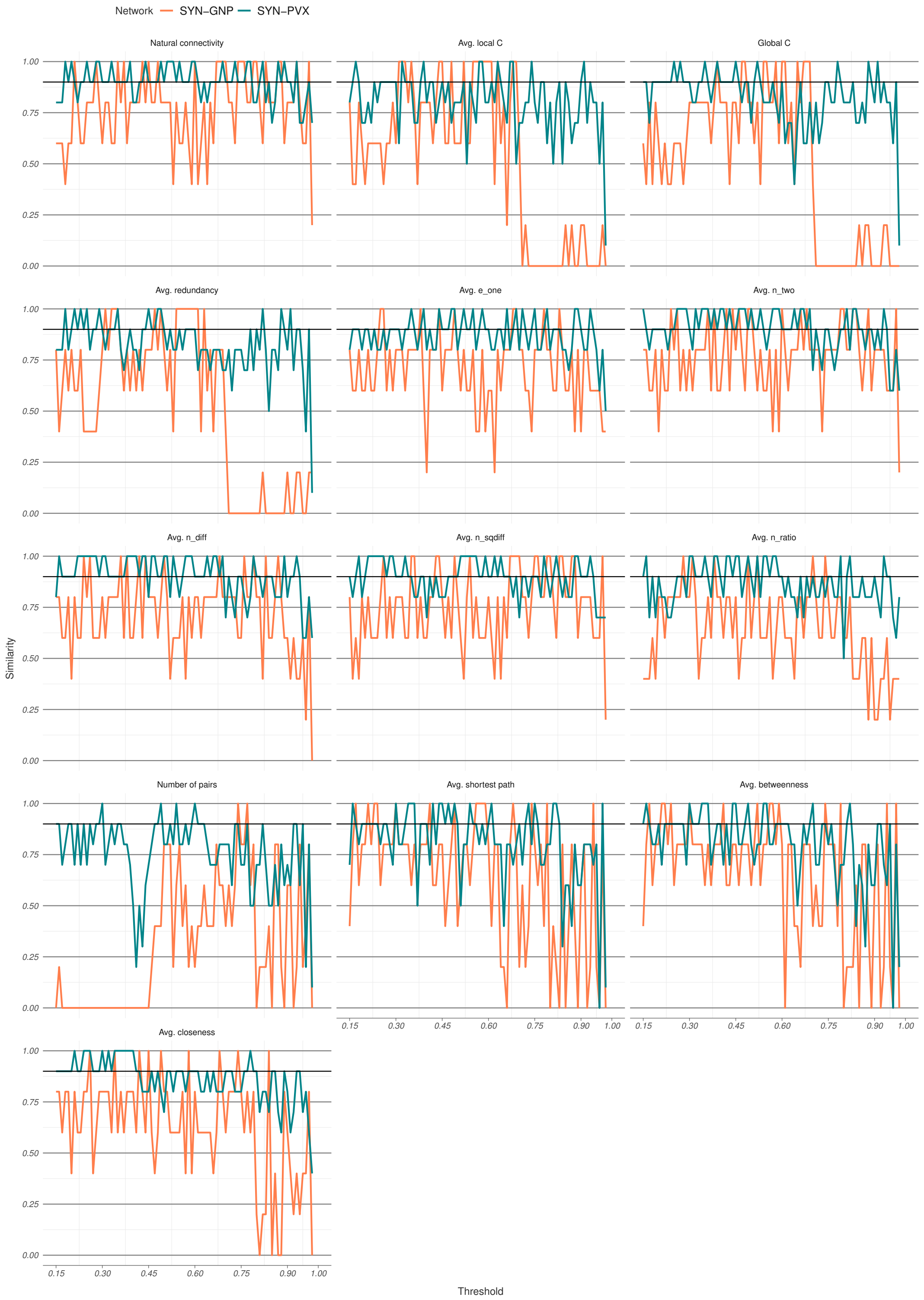

Figure S4: **LOUD metric rank similarity between consecutive thresholds for the synthetic networks.** The SYN-GNP network exhibits lower k-similarity across most metrics and thresholds, as well as generally a wider variability in k-similarity scores.

### 3 Rank identifiability: additional results

| Metric | PVX | ECOLI | YEAST | HPRED | SYN-GNP | SYN-PVX |
|---|---|---|---|---|---|---|
| LOUD Avg. betweenness | 0.22 | 0.07 | 0.21 | 0.35 | 0.05 | 0.51 |
| LOUD Avg. closeness | 0.63 | 0.55 | 0.58 | 0.88 | 0.20 | 0.86 |
| LOUD Avg. Ego1: edges | **0.97** | **0.92** | **0.96** | **0.98** | 0.40 | **0.94** |
| LOUD Avg. local clustering | 0.45 | 0.30 | 0.38 | 0.69 | 0.10 | 0.54 |
| LOUD Avg. Ego2-Ego1: nodes | 0.62 | 0.74 | 0.85 | **0.92** | 0.45 | **0.94** |
| LOUD Avg. Ego1/Ego2: nodes | 0.44 | 0.42 | 0.51 | 0.39 | 0.35 | 0.40 |
| LOUD Avg. $Ego1^2$-Ego2: nodes | **0.94** | 0.86 | 0.87 | **0.97** | 0.40 | **0.94** |
| LOUD Avg. Ego2: nodes | 0.72 | 0.74 | 0.87 | **0.95** | 0.45 | **0.93** |
| LOUD Avg. shortest path | 0.22 | 0.02 | 0.43 | 0.63 | ∼0 | 0.50 |
| LOUD Avg. redundancy | 0.89 | **0.96** | **0.96** | **0.99** | ∼0 | 0.56 |
| Betweenness | 0.68 | 0.73 | 0.74 | **0.93** | 0.30 | 0.85 |
| Closeness | 0.74 | 0.77 | 0.82 | **∼1** | 0.25 | 0.89 |
| Harmonic centrality | 0.82 | 0.77 | 0.81 | **0.97** | 0.35 | **0.91** |
| Degree | 0.86 | 0.87 | 0.89 | **0.99** | 0.40 | **0.95** |
| Ego1: edges | **0.97** | **0.92** | **0.96** | **0.98** | 0.50 | **0.93** |
| LOUD Global clustering | 0.89 | **0.93** | **0.98** | **0.98** | 0.05 | 0.53 |
| Local clustering | 0.25 | 0.22 | 0.20 | 0.46 | ∼0 | 0.36 |
| Ego2-Ego1: nodes | 0.59 | 0.73 | 0.78 | 0.86 | 0.40 | **0.92** |
| Ego1/Ego2: nodes | 0.09 | 0.14 | 0.12 | 0.13 | 0.05 | 0.37 |
| $Ego1^2$-Ego2: nodes | 0.86 | 0.88 | 0.88 | **0.99** | 0.45 | **0.91** |
| Ego2: nodes | 0.68 | 0.74 | 0.79 | **0.96** | 0.45 | **0.92** |
| LOUD Natural connectivity | **0.99** | **0.98** | **0.96** | **∼1** | 0.45 | **0.92** |
| LOUD No connected node pairs | 0.41 | 0.23 | 0.25 | 0.45 | 0.15 | 0.44 |
| PageRank | 0.83 | 0.83 | 0.89 | **∼1** | 0.45 | **0.91** |
| Redundancy | **0.94** | **0.99** | **∼1** | **0.95** | ∼0 | 0.54 |

Table S6: **Identifiability for the 25 metrics across each of the six scored networks.** Values over 0.90 have been highlighted. The SYN-GNP network exhibits generally lower rank identifiability across node metrics compared to the scored PINs. The often higher SYN-PVX identifiability may be due to the random allocation of scores among network edges. This results in edges being deleted homogeneously across the network as the threshold is increased, meaning higher degree nodes are more likely to remain at relatively high degree at different thresholds.

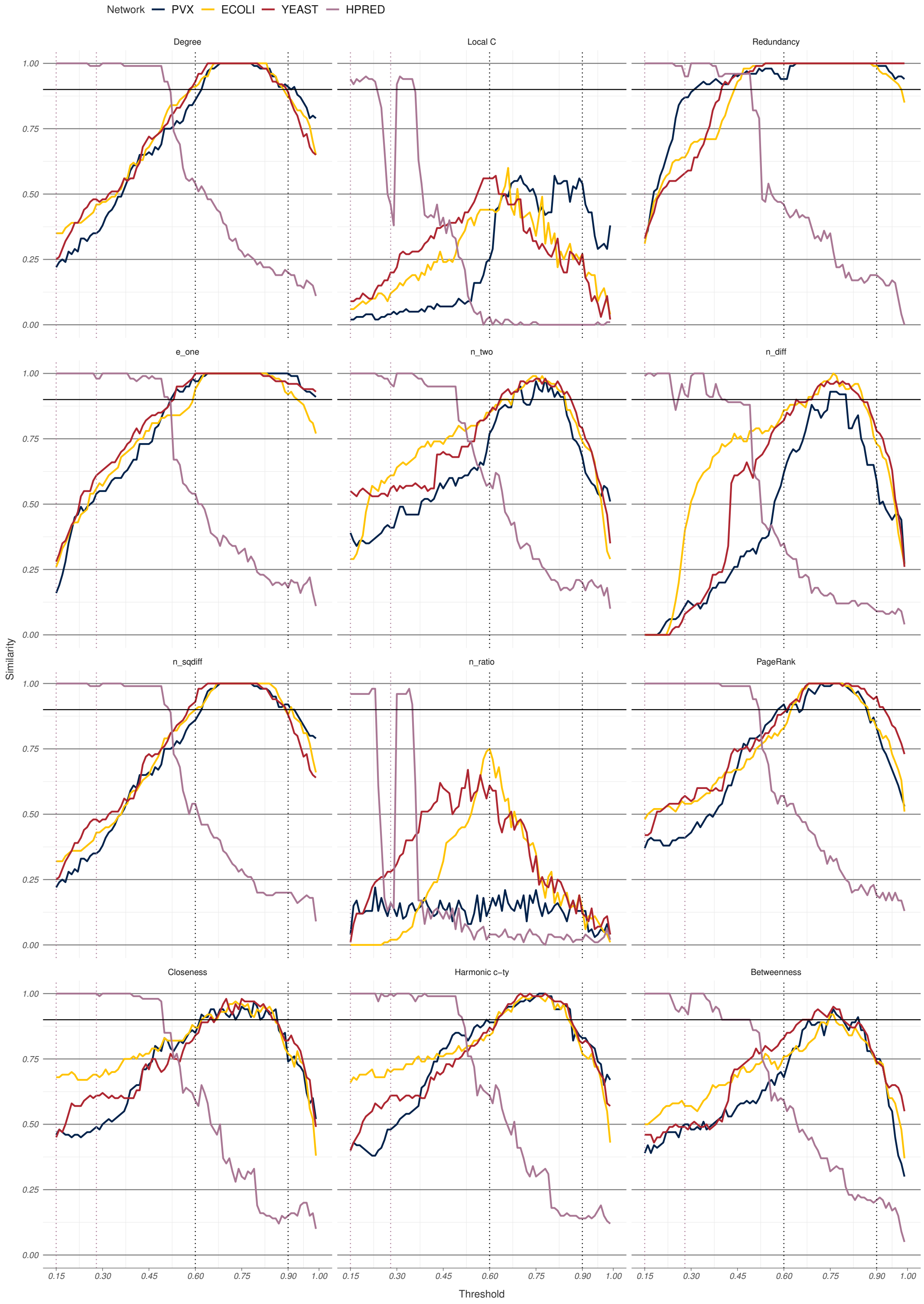

Figure S5: **Standard metric relaxed similarity between thresholded and overall ranks for the four PINs.** The three STRING networks show generally good agreement. The HPRED network, which has been optimised over a different medium-high threshold region (0.15 to 0.28 as opposed to 0.60 to 0.90) behaves significantly differently as the threshold changes.

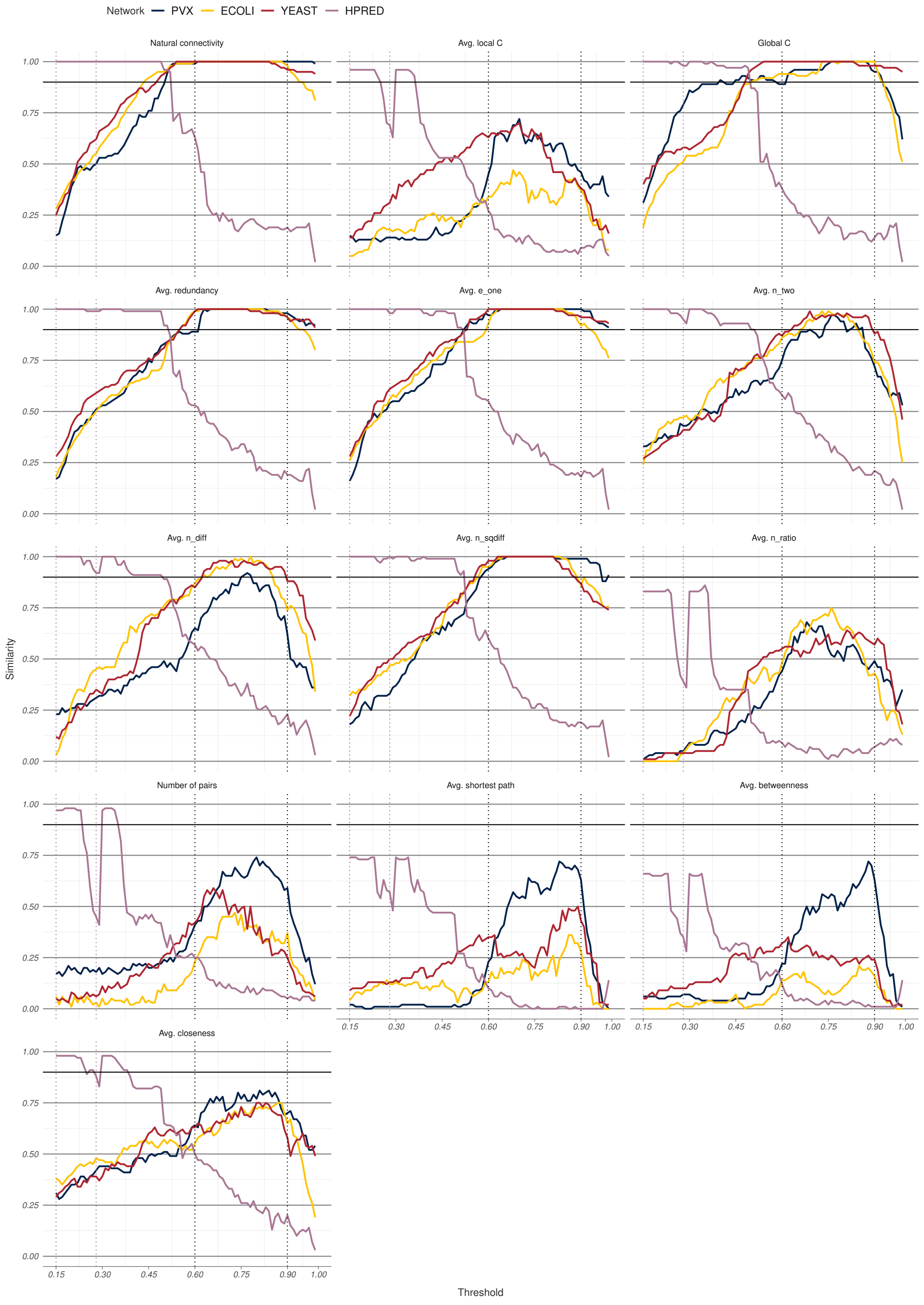

Figure S6: **LOUD metric relaxed rank similarity between between thresholded and overall ranks for the four PINs.** As with the standard metrics (Figure S5), relaxed similarity in the HPRED network behaves differently as a function of the threshold.

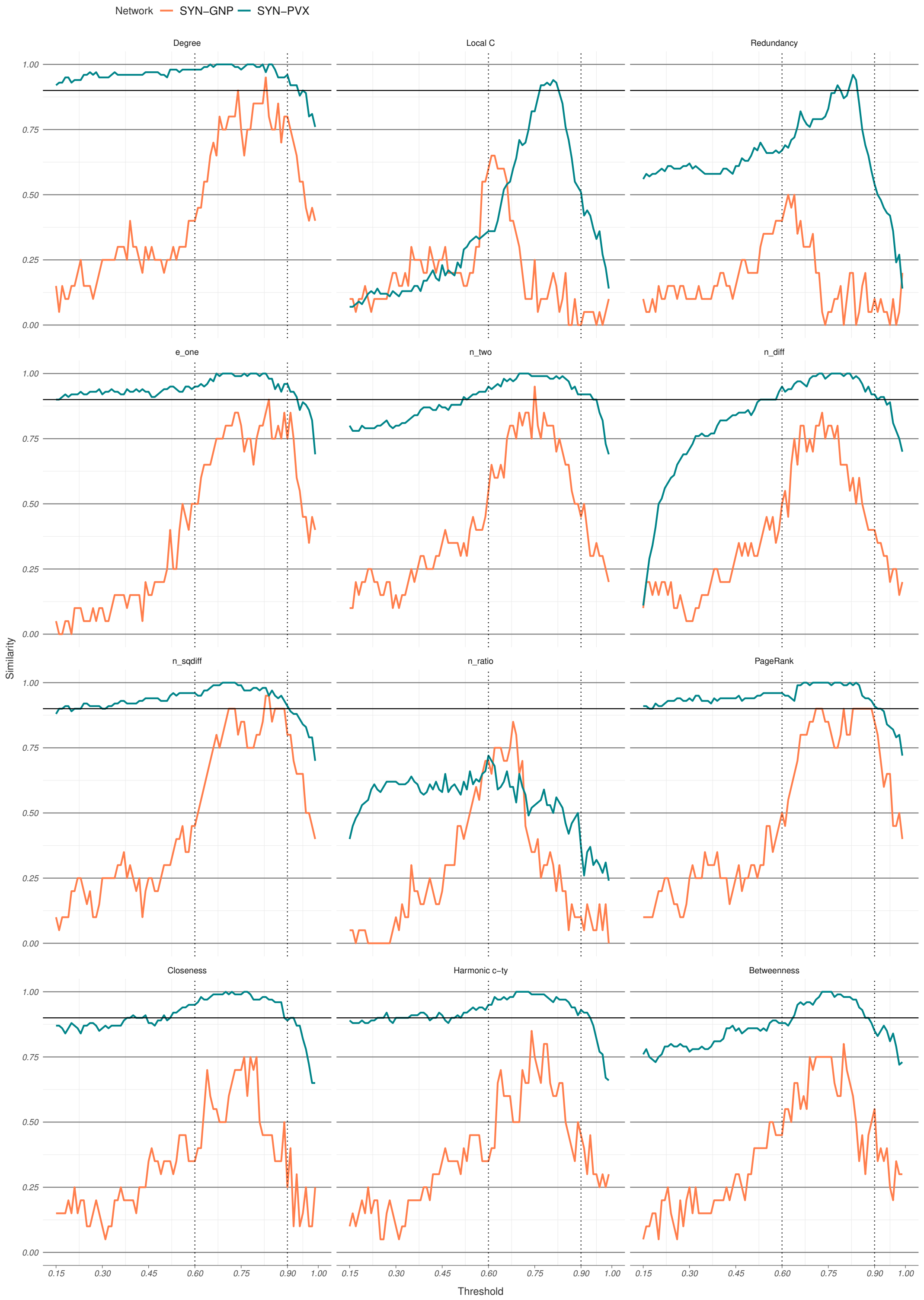

Figure S7: **Standard metric relaxed similarity between thresholded and overall ranks for the synthetic networks.** The SYN-PVX network shows better relaxed rank similarity than the three STRING PINs (Figure S5) overall, while the SYN-GNP network almost never reaches relaxed similarity of 0.90.

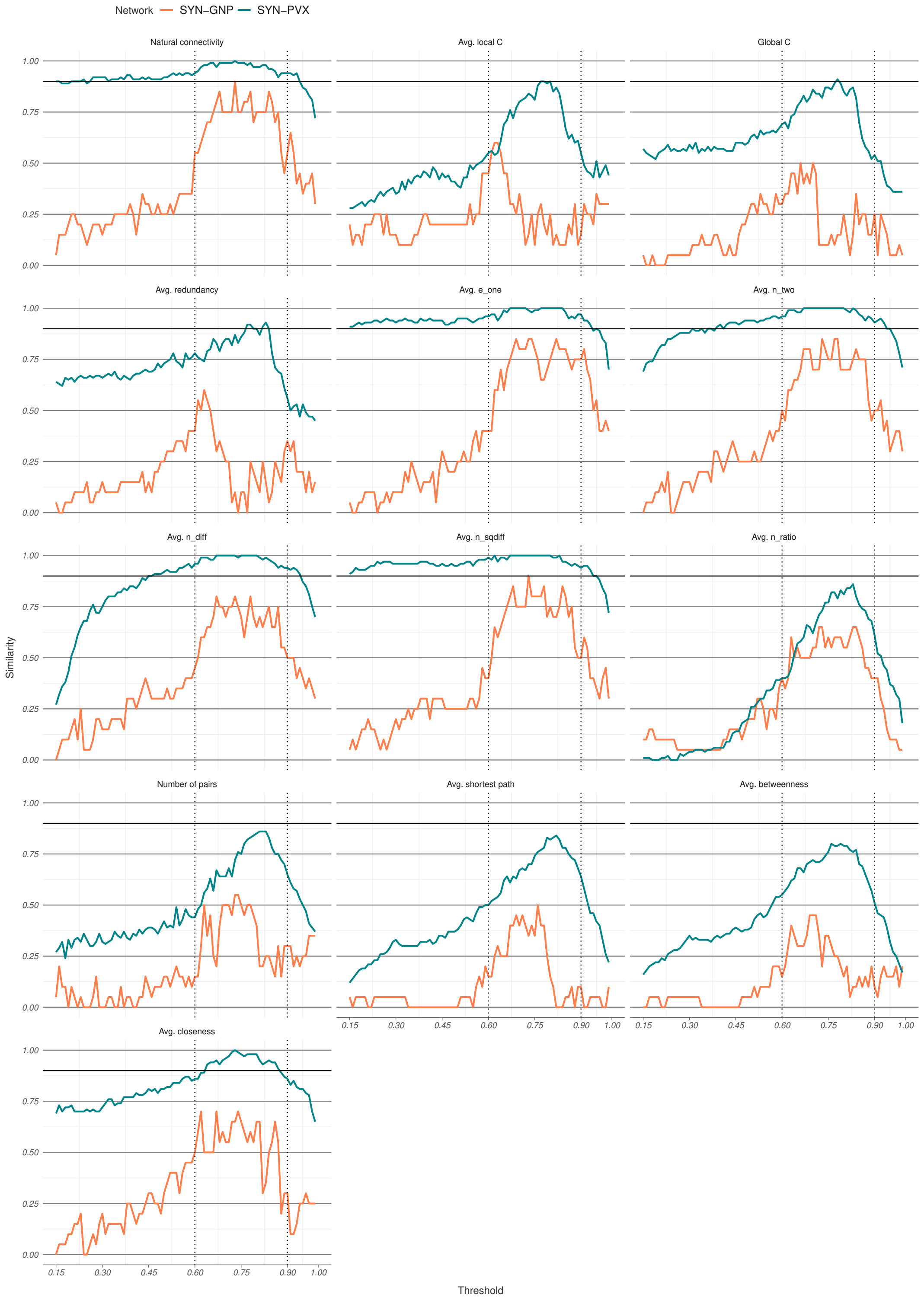

Figure S8: **LOUD metric relaxed rank similarity between between thresholded and overall ranks for the synthetic networks.** The SYN-PVX network shows better relaxed rank similarity than the three STRING PINs (Figure S6) overall, while the SYN-GNP network almost never reaches relaxed similarity of 0.90.

### 4 Rank instability: additional results

| Metric | PVX | ECOLI | YEAST | HPRED | SYN-GNP | SYN-PVX |
|---|---|---|---|---|---|---|
| LOUD Avg. betweenness | 0.09 | 0.22 | 0.12 | 0.02 | 0.44 | 0.07 |
| LOUD Avg. closeness | 0.04 | 0.03 | 0.03 | **0.01** | 0.35 | 0.05 |
| LOUD Avg. Ego1: edges | **0.01** | **∼0** | **∼0** | **∼0** | 0.11 | 0.01 |
| LOUD Avg. local clustering | 0.08 | 0.13 | 0.05 | **0.01** | 0.55 | 0.16 |
| LOUD Avg. Ego2-Ego1: nodes | 0.03 | 0.01 | **0.01** | **∼0** | 0.09 | 0.02 |
| LOUD Avg. Ego1/Ego2: nodes | 0.08 | 0.04 | 0.04 | 0.02 | 0.21 | 0.08 |
| LOUD Avg. $\text{Ego1}^2$-Ego2: nodes | 0.01 | **0.01** | **0.01** | **∼0** | 0.08 | 0.01 |
| LOUD Avg. Ego2: nodes | 0.02 | 0.01 | **0.01** | **∼0** | 0.08 | 0.02 |
| LOUD Avg. shortest path | 0.08 | 0.17 | 0.10 | 0.02 | 0.41 | 0.09 |
| LOUD Avg. redundancy | **0.01** | **∼0** | **∼0** | **∼0** | 0.60 | 0.08 |
| Betweenness | 0.02 | 0.02 | 0.01 | **∼0** | 0.12 | 0.02 |
| Closeness | 0.03 | 0.02 | 0.01 | **∼0** | 0.21 | 0.02 |
| Harmonic centrality | 0.02 | 0.02 | 0.01 | **∼0** | 0.12 | 0.01 |
| Degree | 0.01 | 0.01 | **0.01** | **∼0** | 0.07 | 0.01 |
| Ego1: edges | **0.01** | **∼0** | **∼0** | **∼0** | 0.09 | 0.01 |
| LOUD Global clustering | 0.01 | **∼0** | **0.01** | **∼0** | 0.52 | 0.07 |
| Local clustering | 0.07 | 0.12 | 0.06 | 0.03 | 0.92 | 0.22 |
| Ego2-Ego1: nodes | 0.05 | 0.02 | 0.01 | 0.01 | 0.12 | 0.02 |
| Ego1/Ego2: nodes | 0.46 | 0.27 | 0.15 | 0.05 | 0.38 | 0.31 |
| $\text{Ego1}^2$-Ego2: nodes | 0.01 | 0.01 | **0.01** | **∼0** | 0.08 | 0.01 |
| Ego2: nodes | 0.04 | 0.02 | 0.01 | **∼0** | 0.09 | 0.01 |
| LOUD Natural connectivity | **0.01** | **∼0** | **∼0** | **∼0** | 0.09 | 0.02 |
| LOUD No connected node pairs | 0.25 | 0.53 | 0.54 | 0.09 | 0.55 | 0.11 |
| PageRank | 0.02 | 0.01 | **0.01** | **∼0** | 0.07 | 0.01 |
| Redundancy | 0.01 | **0.01** | **0.01** | **∼0** | 0.96 | 0.07 |

Table S7: **Instability for the 25 metrics across each of the six scored networks.** Values under 0.01 have been highlighted. Values rounded down to 0.01 have not been highlighted. Both synthetic networks tend to have higher instability than the scored PINs across all node metrics.

### 5 STRING network v.9 and v.10

Score distributions for interactions in yeast (*S. cerevisiae*) were compared between STRING v.9 and STRING v.10. The latter was used to build the YEAST network. STRING v.9 contains 777,589 distinct scored interactions, while STRING v.10 contains 939,998 scored interactions. The overlap between the two is 540,162 interactions, or approximately 69% of the older dataset, and 57% of the more recent one. The vast majority of shared interactions were rescored in the update, with approximately 39% of interactions undergoing an absolute change in score above 0.10 and 9% of interactions undergoing an absolute change in score above 0.30. Score distributions are shown in Figure S5.

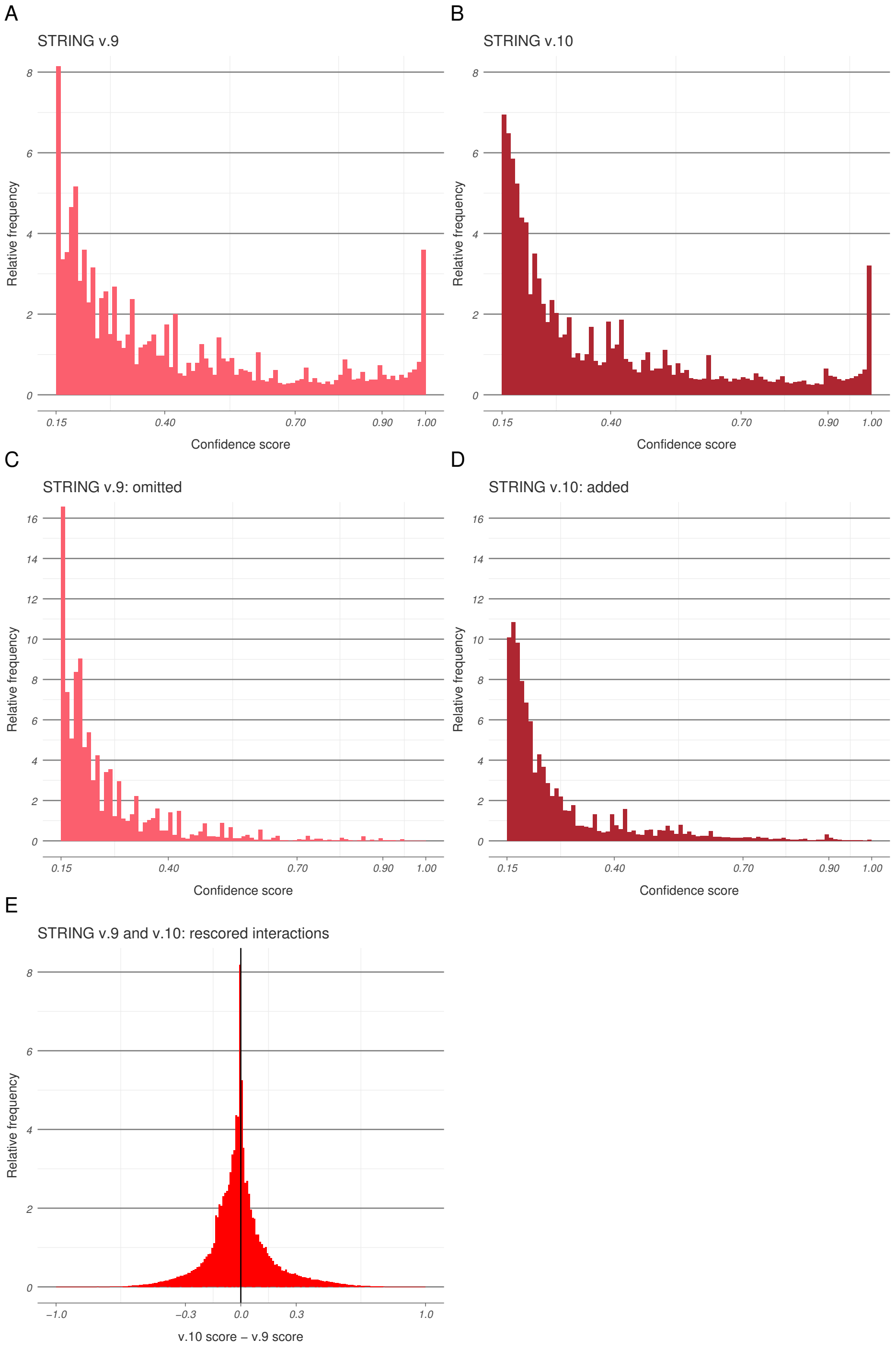

Figure S9: **Confidence score distributions for yeast interactions in STRING v.9 and v.10.** Score distributions for all recorded interactions in v.9 (A) and v.10 (B) are different, although in both cases the majority of interactions have low scores, and a small number of verified interactions have scores close to one. The interactions which appear only in v.9 but were excluded from v.10 (C) have predominantly low scores, although some interactions with high scores were also excluded. Interactions which were added to v.10 but do not appear in the previous version (D) are also predominantly low-scoring. The difference between v.10 and v.9 scores of shared interactions (E) has mean -0.006 and standard deviation 0.168.

### 6 Metric extraction

Node metrics were calculated for all thresholded networks. These metrics can be divided in two groups, which we call "standard" and "leave-one-out difference" (LOUD). Standard metrics describe how a node is embedded in the network. These include well-studied metrics such as local clustering coefficient and node betweenness, as well as metrics based on properties of the node's step-one and step-two ego networks. LOUD metrics aim to quantify the effect of isolating a node in the network. These are global metrics, which were evaluated at a perturbed $G_v^{th}$ for every node $v$. Where possible, a LOUD equivalent was derived for each standard metric.

In order to reduce computational cost, LOUD metrics were calculated only for nodes with degree at least two. LOUD metric values which were not calculated were set to NA (not available) and ranked last in subsequent analysis. Since leaves and isolated nodes are unlikely to be highly ranking with respect to any of the analysed metrics, reducing the calculation in this way should not impact overall analysis.

#### 6.1 Standard metrics

Standard metrics were calculated for each node and at every threshold. These included the degree, local clustering coefficient, redundancy, PageRank centrality, closeness centrality, valued or harmonic centrality, and vertex betweenness. Unless stated otherwise, all metrics were calculated in the conventional way.

*PageRank* was calculated with the default damping factor $d = 0.85$.

*Closeness centrality* was calculated as

$$C(v) = \frac{1}{\sum_{i \neq v} d(v,i)},$$

where the distance between vertices in different connected components of the graph was resolved as $d(u,v) = |V(G)|$.

*Harmonic centrality* is a variation of closeness centrality:

$$H(v) = \sum_{i \neq v} \frac{1}{d(v,i)}.$$

It is defined unambiguously for networks with multiple connected components by taking $1/d(u,v) = 0$ for disconnected nodes.

Basic properties of the step-one and step-two ego networks were also considered. These included

- $e_{one}(v) = |E(ego_1(v))|$, the number of edges in the step-one ego network,
- $n_{two}(v) = |V(ego_2(v))|$, the number of nodes in the step-two ego network,
- $n_{diff}(v) = n_{two}(v) - n_{one}(v)$, the number of nodes within distance exactly two from the node,
- $n_{sqdiff}(v) = n_{one}(v)^2 - n_{two}(v)$, a measure of relative density, and
- $n_{ratio}(v) = n_{one}(v)/n_{two}(v)$, the ratio of neighbourhood sizes.

Above $n_{one}(v) = degree(v) + 1$ is the number of nodes in the step-one ego network. Since this is a linear function of the degree and induces the same ranking, it was not considered as a separate node metric.

#### 6.2 Leave-one-out difference metrics

When calculating LOUD metrics, we considered perturbing the thresholded network $G^{th}$ by isolating each vertex in turn, defining $G_v^{th} = (V(G^{th}), E(G^{th}) \setminus \{(v,i) : i \in V\})$. Then any global metric $f : G^{th} \to \mathbb{R}$ can be re-defined as

$$f_{LOUD}^{th}(v) = f(G^{th}) - f(G_v^{th}).$$

Since only node ranks, rather than exact metric values, were of interest, it was sufficient to calculate $\tilde{f}_{LOUD}^{th}(v) = -f(G_v^{th})$.

Global metrics redefined as leave-one-out included average local clustering coefficient, global clustering coefficient, average redundancy, average path length (i.e. mean geodesic distance), average closeness, average betweenness, number of connected pairs, and natural connectivity, as well as averages of all five standard metrics based on ego networks.

Where possible, a LOUD equivalent was calculated for each standard metric. *Degree* was omitted, since the average degree in the perturbed network results in the same ranking as simply calculating the degree. *PageRank* has no natural global metric extension and was therefore also omitted.

For the *average path length*, only paths between connected vertices were considered. The *average closeness* was calculated as the inverse of the average path length.

*Average betweenness* was calculated using the result in Proposition 1 as

$$\overline{btw} \propto \sum_{w} btw(w) = N_2(\bar{l} - 1).$$

In the above, the *number of connected pairs* $N_2$ is the number of pairs of nodes within the same component, and $\bar{l}$ is the average path length.

**Proposition 1.** *Let $G^{th} = (V, E^{th})$ be a simple, undirected graph on $|V| = n$ nodes. Let $btw(w)$ be the betweenness centrality for the node $w$, $N_2$ be the number of connected node pairs, and $\bar{l}$ be the mean geodesic distance of $G^{th}$. Then*

$$\sum_{w} btw(w) = N_2(\bar{l} - 1).$$

*Proof.* For a triplet of connected nodes $u, v, w \in V$, define the following:

- $l_{uv}$ = the shortest path length between $u$ and $v$,
- $\sigma_{uv}$ = the number of distinct shortest paths between $u$ and $v$,
- $\sigma_{uv}^{w}$ = the number of distinct shortest paths between $u$ and $v$ which pass through $w$.

Then betweenness is defined as

$$btw(w) = \sum_{u,v \neq w} \frac{\sigma_{uv}^{w}}{\sigma_{uv}},$$

where the sum is over node pairs $(u, v)$ in the same component as $w$. Summing over $w$ and swapping the sums,

$$\sum_{w} btw(w) = \sum_{u,v} \sum_{w \neq u,v} \frac{\sigma_{uv}^{w}}{\sigma_{uv}} = \sum_{u,v} \frac{1}{\sigma_{uv}} \sum_{w \neq u,v} \sigma_{uv}^{w}.$$

If $u$, $v$ and $w$ are in the same component, in $\sum_{w} \sigma_{uv}^{w}$, each path of length $l_{uv}$ contributes $l_{uv} - 1$, as there are $l_{uv} - 1$ intermediate nodes on such a path. Therefore, $\sum_{w} \sigma_{uv}^{w} = \sigma_{uv}(l_{uv} - 1)$ and so

$$\sum_{w} btw(w) = \sum_{u,v} \frac{1}{\sigma_{uv}} \sigma_{uv}(l_{uv} - 1) = \sum_{u,v} (l_{uv} - 1) = N_2\bar{l} - N_2 = N_2(\bar{l} - 1).$$

□

*Natural connectivity* is a metric of global network robustness. It is calculated using the spectrum of the adjacency matrix of $G^{th}$. If the eigenvalues of the adjacency are $\lambda_i, i \in 1, \ldots, n$, then the natural connectivity of $G^{th}$ is

$$N(G^{th}) = log\Big(\frac{1}{n} \sum_{i=1}^{n} e^{\lambda_i}\Big).$$

The complete set of the 25 metrics used can be found in Table S8.

| Standard | LOUD |
|---|---|
| Degree | |
| Local clustering | Average local clustering |
| | Global clustering |
| Redundancy | Average redundancy |
| PageRank | |
| Closeness | Average closeness |
| Harmonic centrality | Average path length |
| | Number of connected pairs |
| Betweenness | Average betweenness |
| | Natural connectivity |
| $e_{one}(v)$ | Average $e_{one}(v)$ |
| $n_{two}(v)$ | Average $n_{two}(v)$ |
| $n_{diff}(v)$ | Average $n_{diff}(v)$ |
| $n_{sqdiff}(v)$ | Average $n_{sqdiff}(v)$ |
| $n_{ratio}(v)$ | Average $n_{ratio}(v)$ |

Table S8: **The complete set of twenty-five standard and leave-one-out metrics.** Metrics in the same row of the table aim to capture similar properties in a standard and LOUD form.

### 7 Worked example

In order to assess rank robustness for a node metric in a particular scored network, first the metric is calculated at all relevant threshold, and then threshold node rankings are extracted. This is enough to calculate rank continuity for the metric. Overall rankings are then calculated, and used together with the threshold rankings to calculate rank identifiability and rank instability. The following is an example of how the rank robustness of node degree is measured in the PVX network.

#### 7.1 Metric extraction and ranking

Consider a random subset of twenty nodes in the PVX network (Table S9), the degrees of which are observed at the start and end of the medium-high confidence region, i.e. at thresholds 0.60 and 0.90.

The majority of nodes in the network have low degree, and may be isolated at higher thresholds—for example, nine of the twenty nodes have degree 0 at threshold 0.60, and fourteen have degree 0 at threshold 0.90. Since higher thresholds correspond to edge deletion in the network, degree behaves as a decreasing function of the score threshold.

After node metrics, such as degree, are calculated, nodes are ranked at every threshold. Higher ranks correspond to higher metric values. Therefore, PVX_000575, which has degree 1 at threshold 0.60, has a higher rank (1174) than any of the isolated nodes at that threshold (between 201 and 979 for the nodes in Table S9). Ties are resolved at random, so when PVX_000575 becomes isolated at threshold 0.90, its rank is no longer necessarily higher than that of other isolated nodes (1576 when ranks for isolated nodes in the table range between 445 and 1846).

Overall ranks are calculated by ranking the average ranks within the medium-high confidence region, from 0.60 to 0.90. Due to ties and inhomogeneous score placement across network edges, node ranks do not change monotonically with the threshold, even if the metrics they are based on do. Overall ranks may therefore be lower than they are at either end of the region (e.g. PVX_000575), they may be higher (e.g. PVX_091810), or they may fall between the two (e.g. PVX_000775).

#### 7.2 Rank robustness

*Rank continuity* is based on the amount of overlap between the sets of highest ranking nodes in consecutive thresholds. First, the top $n = Nk$ highest degree nodes are identified at each threshold, for values of $k$ between 0.001 and 0.05 at 0.001 intervals. In the above, $N$ is the total number of nodes (3255 in the PVX network). Then these sets are compared between consecutive thresholds, and their overlap is recorded (see Table S10 for sample values). Finally, the rank continuity score is calculated as the fraction of times the recorded overlap is over 90%, i.e. the fraction of times $M_k > 0.90$. All analysed values of $k$ and all thresholds in the medium-high confidence region are included in calculating a single rank continuity score.

| name | Degree (0.60) | Degree (0.90) | Rank (0.60) | Rank (0.90) | Rank (overall) |
|---|---|---|---|---|---|
| **PVX_000575** | 1 | 0 | 1174 | 1576 | 1073 |
| **PVX_000775** | 38 | 9 | 2371 | 2692 | 2416 |
| PVX_001065 | 0 | 0 | 979 | 1846 | 292 |
| PVX_002905 | 0 | 0 | 835 | 670 | 614 |
| PVX_080050 | 21 | 6 | 2131 | 2507 | 2241 |
| PVX_089440 | 0 | 0 | 585 | 790 | 456 |
| PVX_090015 | 23 | 6 | 2179 | 2476 | 2319 |
| **PVX_091810** | 309 | 90 | 3247 | 3178 | 3253 |
| PVX_094970 | 8 | 0 | 1744 | 854 | 1707 |
| PVX_098000 | 12 | 0 | 1879 | 1147 | 1746 |
| PVX_098940 | 2 | 0 | 1384 | 1609 | 1266 |
| PVX_100565 | 21 | 3 | 2149 | 2284 | 2243 |
| PVX_101285 | 104 | 15 | 2857 | 2842 | 2867 |
| PVX_113865 | 0 | 0 | 576 | 671 | 17 |
| PVX_114885 | 0 | 0 | 646 | 445 | 1119 |
| PVX_117400 | 0 | 0 | 911 | 1777 | 348 |
| PVX_117550 | 0 | 0 | 201 | 1589 | 1043 |
| PVX_117570 | 0 | 0 | 418 | 1153 | 291 |
| PVX_122590 | 1 | 0 | 1291 | 1449 | 1461 |
| PVX_123750 | 0 | 0 | 310 | 1182 | 201 |

Table S9: **Degrees and the rankings induced by them for 20 random nodes in the PVX network.** Highlighted nodes are firther discussed in the text.

| Thresholds | Fraction $k$ of interest | Number of nodes $n = Nk$ | Overlap | Score $M_k$ |
|---|---|---|---|---|
| 0.60 to 0.61 | 0.01 | 32 | 29 | 0.9063 |
| 0.60 to 0.61 | 0.02 | 65 | 64 | 0.9846 |
| 0.75 to 0.76 | 0.01 | 32 | 30 | 0.9375 |
| 0.75 to 0.76 | 0.02 | 65 | 61 | 0.9385 |
| 0.90 to 0.91 | 0.01 | 32 | 30 | 0.9375 |
| 0.90 to 0.91 | 0.02 | 65 | 62 | 0.9538 |

Table S10: **Rank overlap score, i.e. k-similarity, for degree in the PVX network. Selected thresholds and values of $k$ are shown.** All shown values of $M_k$ are over 0.90. If rank continuity was calculated based only on the data in this table, degree would have rank continuity 1 in the PVX network.

| Threshold | 0.60 | 0.65 | 0.70 | 0.75 | 0.80 | 0.85 | 0.90 |
|---|---|---|---|---|---|---|---|
| $M_k^\alpha$ | 0.86 | 0.98 | 1.00 | 1.00 | 0.99 | 0.95 | 0.91 |

Table S11: **Relaxed k-similarity between overall ranks and ranks at selected thresholds, based on degree in the PVX network.** The lowest shown value for $M_k^\alpha$ is $M_k^\alpha = 0.86$ for confidence score threshold 0.60. This is also the lowest value found in the entire medium-high confidence region, which is why degree has rank identifiability 0.86 for the PVX network (see Table S6).

| Name | Rank range | Relative rank range |
|---|---|---|
| PVX_080245 | 23 | 0.00707 |
| PVX_084620 | 81 | 0.02488 |
| PVX_085275 | 20 | 0.00614 |
| PVX_091810 | 74 | 0.02273 |
| PVX_096265 | 31 | 0.00952 |
| PVX_111040 | 79 | 0.02427 |
| PVX_115340 | 48 | 0.01475 |
| PVX_117170 | 24 | 0.00737 |
| PVX_117395 | 50 | 0.01536 |
| PVX_117925 | 32 | 0.00983 |

Table S12: **Rank ranges for the top 10 highest degree nodes in the PVX network.** Rank instability is the average relative rank range, and if only the top 10 nodes were considered it would be 0.014.

Calculating the *rank identifiability* for degree in the PVX network is based on a comparison between the highest degree nodes at different thresholds and the highest degree nodes "overall", i.e. the nodes with highest overall ranks. Rather than varying $n = kN$, the number of nodes is fixed at $n = 100$. First, the set of top 100 overall ranking nodes is identified. Then their ranks at different thresholds of interest are queried (i.e. for all thresholds in the medium-high confidence region). For each threshold, the number of these overall top nodes which are also in the top 150 for the particular threshold, is calculated. When scaled, this is $\alpha$-relaxed k-similarity $M_k^\alpha$ between overall and threshold ranks, with $\alpha = 1.5$ and $k = 100/N$. Sample values can be found in Table S11. Rank identifiability is calculated as the minimum observed $M_k^\alpha$ across the threshold region. Rank identifiability is the fraction of the overall top 100 nodes which are guaranteed to be identified among the top 150, no matter which threshold is chosen.

Finally, *rank instability* provides another comparison between overall ranks and ranks obtained at different thresholds. First the top 1% of overall degree nodes are identified (the top 32 in the PVX network). Then, their ranks are tracked across the medium-high confidence region and the range of ranks attained for each node is recorded (i.e. the difference between the node's highest recorded and lowest recorded ranks). Table S12 contains data for the top 10 highest degree nodes. Ranges are scaled relative to the number of nodes in the network, and rank instability is calculated as the average relative rank range among these top nodes.
